# Genome-wide profiling of microRNAs and prediction of mRNA targets in 17 bovine tissues

**DOI:** 10.1101/574954

**Authors:** Min Wang, Amanda J Chamberlain, Claire P Prowse-Wilkins, Christy J Vander Jagt, Timothy P Hancock, Jennie E Pryce, Benjamin G Cocks, Mike E Goddard, Benjamin J Hayes

## Abstract

MicroRNAs regulate many eukaryotic biological processes in a temporal- and spatial-specific manner. Yet in cattle it is not fully known which microRNAs are expressed in each tissue, which genes they regulate, or which sites a given microRNA bind to within messenger RNAs. An improved annotation of tissue-specific microRNA network may in the future assist with the identification of causal variants affecting complex traits. Here, we report findings from analysing short RNA sequence from 17 tissues from a single lactating dairy cow. Using miRDeep2, we identified 699 expressed mature microRNA sequences. Using TargetScan, known (60%) and novel (40%) microRNAs were predicted to interact with 780,481 sites in bovine messenger RNAs homologous with human. Putative interactions between microRNA families and targets were significantly enriched for interactions from previous experimental and computational identification. Characterizing features of microRNAs and targets, we showed that (1) mature microRNAs derived from different arms of the same precursor targeted different genes in different tissues; (2) miRNA target sites preferentially occurred within gene regions marked with active histone modification; (3) variants within microRNAs and targets had lower allele frequencies than variants across the genome, as identified from 65 million whole genome sequence variants; (4) no significant correlation was found between the abundance of microRNAs and messenger RNAs differentially expressed in the same tissue; (5) microRNAs and target sites weren’t significantly associated with allelic imbalance of gene targets. This study contributes to the goals of Functional Annotation of Animal Genomes consortium to improve the annotation of genomes of domestic animals.

## Introduction

Micro RNAs (miRNAs) are a class of single-stranded, short-length (typically 19-24nt), endogenous, non-coding RNAs (ncRNAs) that are involved in almost all biological processes, including development, differentiation, immunity, reproduction and longevity (Kloosterman and Plasterk 2006; Hasuwa et al. 2013; Renthal et al. 2013; Li and Belmonte 2015; Mehta and Baltimore 2016; Wang et al. 2016a; Cowled et al. 2017; Ioannidis and Donadeu 2017; Bartel 2018). Close to three decades of miRNA studies have revealed that miRNAs have broadly conserved biogenesis and conserved target sites at the three prime untranslated regions (3’UTRs) of the messenger RNAs (mRNAs) in eukaryotes (Bartel 2018). Mutations in miRNA target sites have been linked to numerous complex phenotypes in humans, livestock and plants (Mallory et al. 2004; Abelson et al. 2005; Clop et al. 2006; Kloosterman and Plasterk 2006; Leung and Sharp 2010). For example, in humans a single base deletion in the binding sites of *hsa-miR-189* within the 3’UTR of the *SLITRK1* mRNA was shown to be associated with Tourette’s syndrome (Abelson et al. 2005). In Texel sheep, a single-base-nucleotide substitution had created a binding site for the *miR-1/miR-206* family within the 3’UTR of the *GDF8* mRNA, leading to translational repression of the *GDF8* transcript and ‘double muscling’ (Clop et al. 2006). In Arabidopsis, the *GGU* (Gly) *to GAU* (Asp) single-base-nucleotide substitution within the *PHABULOSA* mRNA, which was associated with the development of stunted leaves, was within the binding site of the *miR165/166* family. When a synonymous single-base-nucleotide substitution *GGA* (Gly) was introduced, which didn’t change the protein sequence but disrupted the miRNA binding site, the same stunted leaf phenotype was observed. This confirmed the disruption of miRNA binding site, not the change of PHABULOSA protein sequence, had caused the stunted leaf development (Mallory et al. 2004).

In cattle the spatial- and temporal-interactions between miRNAs and mRNAs are not yet fully understood or characterised. An improved annotation of tissue-specific miRNA interaction networks will provide opportunities to better examine the effects of mutations in miRNA genes and target sites in the bovine genome, particularly in the era of large collections of whole genome sequence (WGS) variants.

In animals, the miRNA gene is transcribed by RNA polymerase II in the nucleus to produce primary miRNA before being processed into precursor miRNA that consists of a hairpin structure (Lee et al. 2004; Morlando et al. 2008; Pawlicki and Steitz 2008; Nojima et al. 2015). The precursor miRNA is exported to cytosol where the stem loop of the hairpin is cleaved to release the duplex. The duplex consists of two RNA strands that are partially reverse-complementarily matched. One strand will become the dominant functional product (i.e. the mature miRNA) and be incorporated into the Argonaute protein as part of the RNA-induced silencing complex to direct post-transcriptional repression of mRNA targets. The other strand will become passenger sequence (i.e. the star miRNA) which is usually discarded and degraded (Bartel 2018). In cytosol, the mature miRNA is mostly known to bind to the 3’UTR of target mRNA to inhibit translation (Bartel 2018). A contiguous 6-nucleotide Watson-Crick base pairing is required between the mRNA target sequence and the miRNA seed sequence, which is the 2^nd^-7^th^ nucleotides at the 5’-end of the mature miRNA. When a target-seed match is extended to 1^st^ or 8^th^ nucleotide or both nucleotides, the repression effect is higher. Other features including sequence context at the 3’UTRs and open reading frame (ORF) have been shown to affect the repression efficacy (Grimson et al. 2007; Friedman et al. 2009; Garcia et al. 2011; Agarwal et al. 2015). Apart from targeting the 3’UTR, miRNAs have also been shown to target the coding regions (CDS), 5’UTRs and introns in the mRNA (Schnall-Levin et al. 2010; Meng et al. 2013; Li et al. 2016).

Experimental and computational methods have been developed to identify miRNA genes and target sites (Krek et al. 2005; Licatalosi et al. 2008; Chi et al. 2009; Betel et al. 2010; Hafner et al. 2010; Zisoulis et al. 2010; Friedländer et al. 2012; Reczko et al. 2012; Helwak et al. 2013; Grosswendt et al. 2014; Agarwal et al. 2015; Paicu et al. 2017). Particularly with recent advances in high-throughput sequencing, more lowly-abundant miRNAs, which are less conserved than the highly-abundant miRNAs, have been detected with unprecedented sensitivity (Friedländer et al. 2012; Londin et al. 2015). This has already led to the discovery of many novel miRNAs in humans and other mammals (Li et al. 2011; Londin et al. 2015; Penso-Dolfin et al. 2016; Wake et al. 2016; Ji et al. 2017; Wong et al. 2017). In bovine, profiling expressed miRNAs using high-throughput sequencing has been performed in a number of tissues and developmental stages (Gu et al. 2007; Jin et al. 2009; Zhixiong et al. 2014). High-throughput sequencing of RNA isolated by crosslinking and immunoprecipitation (HITS-Seq, also known as CLIP-Seq) was recently used to identify the mRNA sequences that were bound by the miRNA-directed Argonaute protein in bovine kidney cells on a genome-wide scale (Scheel et al. 2017). However, because miRNAs have well-defined temporal- and spatial-expression patterns (Guo et al. 2014), we reasoned that many more miRNAs and their target sites are present in cattle and can be identified through analysing additional samples representing more diverse tissue types.

In this study, we present a genome-wide identification, quantification and differential expression of known and novel miRNAs in 17 tissues from a single lactating dairy cow. We predicted the target sites of all expressed miRNAs in bovine mRNA transcripts and demonstrated an enrichment of putative interactions between miRNA families and cognate targets in interactions from previous experimental and computational identification. We summarized features of expressed miRNAs and target sites. We showed a preferential localisation of miRNA target sites at active bovine enhancer regions. Using 65 million WGS variants from the 1000 Bull Genomes Project (Daetwyler et al. 2014) (*Bos Taurus* Run 6), we found rare sequence variants were significantly enriched within miRNA genes and target sites compared with the entire genome. But we didn’t find the expression levels of mature miRNAs significantly associated with allelic imbalance within miRNA target sites, nor did we find heterozygous sites within miRNA target sites significantly contributed to allelic imbalance within exons of the same target genes. Our study provides an updated annotation of bovine miRNA-mRNA regulatory networks that in the future may aid in the identification of causal variants affecting complex traits.

## Results

### Samples, RNA sequencing and alignment

The miRNA profiles were generated by sequencing short RNA libraries prepared in duplicate from adrenal gland, black skin, brain caudal lobe, brain cerebellum, heart, intestinal lymph node, kidney, leg muscle (semimembranosus), liver, lung, mammary gland, ovary, spleen, thymus, thyroid, tongue, and white skin, which were collected from a first-lactation dairy cow as part of another study (Chamberlain et al. 2015). Over 800 million paired-end reads, 5.8-17.8 million reads per library, were generated (Supplemental Table S1). Two trimmers, cutadapt (Martin 2011) in combination with sickle (Joshi and Fass 2011) and trimmomatic (Bolger et al. 2014), were used to compare the efficacy of trimming. Although no adapter contamination was detected by FastQC (Andrews 2010) in any trimmed reads, a much larger number of reads were retained by cutadapt and sickle (83.25-95.88%) than trimmomatic (14.72-60.51%) (Supplemental Table S1).

In order to assess whether adapter residues remained in trimmed libraries, and to identify the best aligner for short RNA sequence reads, reads output from both trimmers were respectively mapped with five aligners: BWA (Li and Durbin 2009), bowtie (Langmead 2010), bowtie2 (Langmead and Salzberg 2012), STAR (Dobin et al. 2013) and HISAT2 (Kim et al. 2015). Across all aligners, sequence reads that were trimmed by the single-end trimmer (i.e. cutadapt and sickle) were aligned to bovine reference genome with much larger gaps than those that were trimmed by the paired-end trimmer (i.e. trimmomatic). Since trimmomatic has the advantage over cutadapt and sickle to utilise paired-end information to identify adapter residues, this observation indicated that adapter residues remained in the single-end trimmed reads even though FastQC (Andrews 2010) could not detect them. Across all libraries, only when short RNA sequence reads were trimmed by trimmomatic and aligned to bovine reference genome using bowtie2, the averaged inferred insert size matched with the insert size from experimental design (10-50nt; Table 1). Bowtie2 was also one of the best two aligners in returning more paired and mapped reads, more exact matched reads, more inward oriented pairs and less low mapping quality reads (Supplemental Table S1). Read depths varied between tissues as expected. Tongue had the largest proportion of mapped and paired reads (98.19%), and black skin had the lowest ratio (90.45%; Supplemental Table S1). All subsequent analyses were progressed with reads processed by trimmomatic and bowtie2.

**Table 1.**
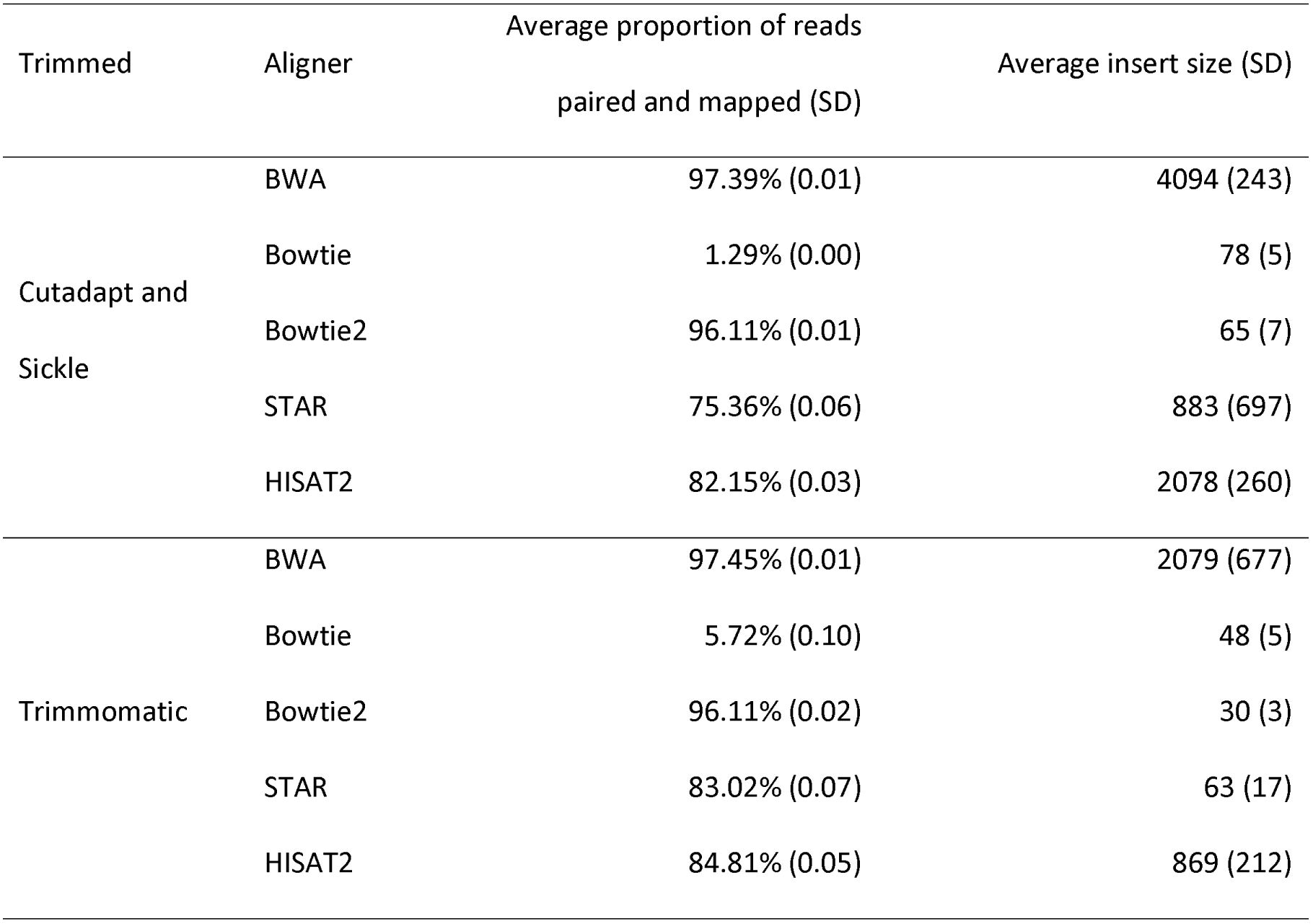
Comparison of Trimmers and Aligners for Short RNA Sequence Reads. For the two trimmers and each of five aligners used, this table presents the average proportions of trimmed reads that were paired and mapped to bovine reference genome and the standard deviation (SD). Also presented is the inferred insert sizes averaged across all libraries.

### Identification of expressed known and novel miRNAs

A total of 13,807 candidate precursor miRNAs genomic coordinates along with their cognate mature and star miRNA sequences from 34 libraries (17 tissues x 2 technical replicates) were identified by miRDeep2 (Friedländer et al. 2012) (Supplemental Table S2). Of those 7,862 passed filtering (false discovery rate ≤0.1 for novel miRNAs), which included 655 unique precursor sequences (44-149nt) that were expressed from 657 genomic coordinates, 699 unique mature miRNA sequences (18-25nt) from 722 genomic coordinates, and 736 unique star miRNA sequences (13-28nt) from 748 genomic coordinates. The inconsistency between the number of unique sequences (precursor, mature or star) and the number of unique genomic locations were due to identical miRNA sequences expressing from >1 distinct genomic location. For example, a novel precursor sequence was expressed from multiple genomic coordinates including location 1: *chr25:32389474-32389547(+)*, location 2: *chr25:32411299-32411372(+)* and location 3: *chr25:32395491-32395564(+)*. All three locations were expressed in ovary; whereas in adrenal gland only location 1 and 2 were expressed. In all three locations, many mature reads (read count: 544-2521) were observed from each tissue. In ovary, a small amount of loop and star miRNAs were observed. Because the loop and star miRNAs degrade quickly, the detection of these sequences along with the high levels of mature sequences increased the confidence of a true detection (Friedländer et al. 2012) (accuracy: 73-79%). Other examples of identical known miRNA sequences (e.g. *bta-miR-7-5p*) or novel miRNA sequences (e.g. *ACUAUACAACCUACUACCUCA*) that were expressed from different genomic locations (within the same chromosome or between chromosomes) in different tissues are also observed (Supplemental Table S2).

Sixty percent of the miRNAs identified were nearly identical to bovine, goat or sheep miRNAs that were annotated in miRBase (Kozomara and Griffiths-Jones 2014) (version 22), and the remaining 40% were novel (Supplemental Table S2). miRDeep2 (Friedländer et al. 2008; Friedländer et al. 2012) classified known and novel miRNAs by comparing our expressed precursor miRNA sequence with the precursor miRNA sequences in miRBase (Kozomara and Griffiths-Jones 2014) (version 22). If an expressed mature miRNA has never been reported to miRBase but is excised from a precursor miRNA in miRBase, the mature miRNA sequence is still classified as ‘known’ (e.g. *bta-miR-26c-5p;* Supplemental Table S2). Because such cases were rare, in this study we used the miRDeep2 classification of known and novel miRNAs.

Known miRNAs were more abundant than novel miRNAs (Figure 1A). Close to 48.19% of the novel mature miRNAs overlapped with the antisense strand of known miRNAs, and these ‘known-antisense’ novel miRNAs were more abundant than the rest of the novel miRNAs (Figure 1B). Known and novel mature miRNAs were more abundant in brain cerebellum than all other tissues (Figure 1C). Most mature miRNAs were expressed in either only one tissue (where >67% were novel) or all 17 tissues (where >98% were known; Figure 1D). Across all tissues, most miRNAs were expressed in clusters within 3,000nt of each other (Figure 2). Many known and novel mature miRNAs were identified on chromosome 21, X and 19 (Figure 1E). No known nor novel miRNAs were identified on the bovine mitochondrial chromosome, even though miRNAs have previously been identified in the mitochondria of liver cells from humans, mice and rats (Kren et al. 2009). This could be because mitochondria have a unique population of miRNAs, and required the isolation of mitochondria from tissue samples to enable RNAs to be extracted from mitochondria as opposed from total cellular content (Kren et al. 2009).

**Figure 1.**
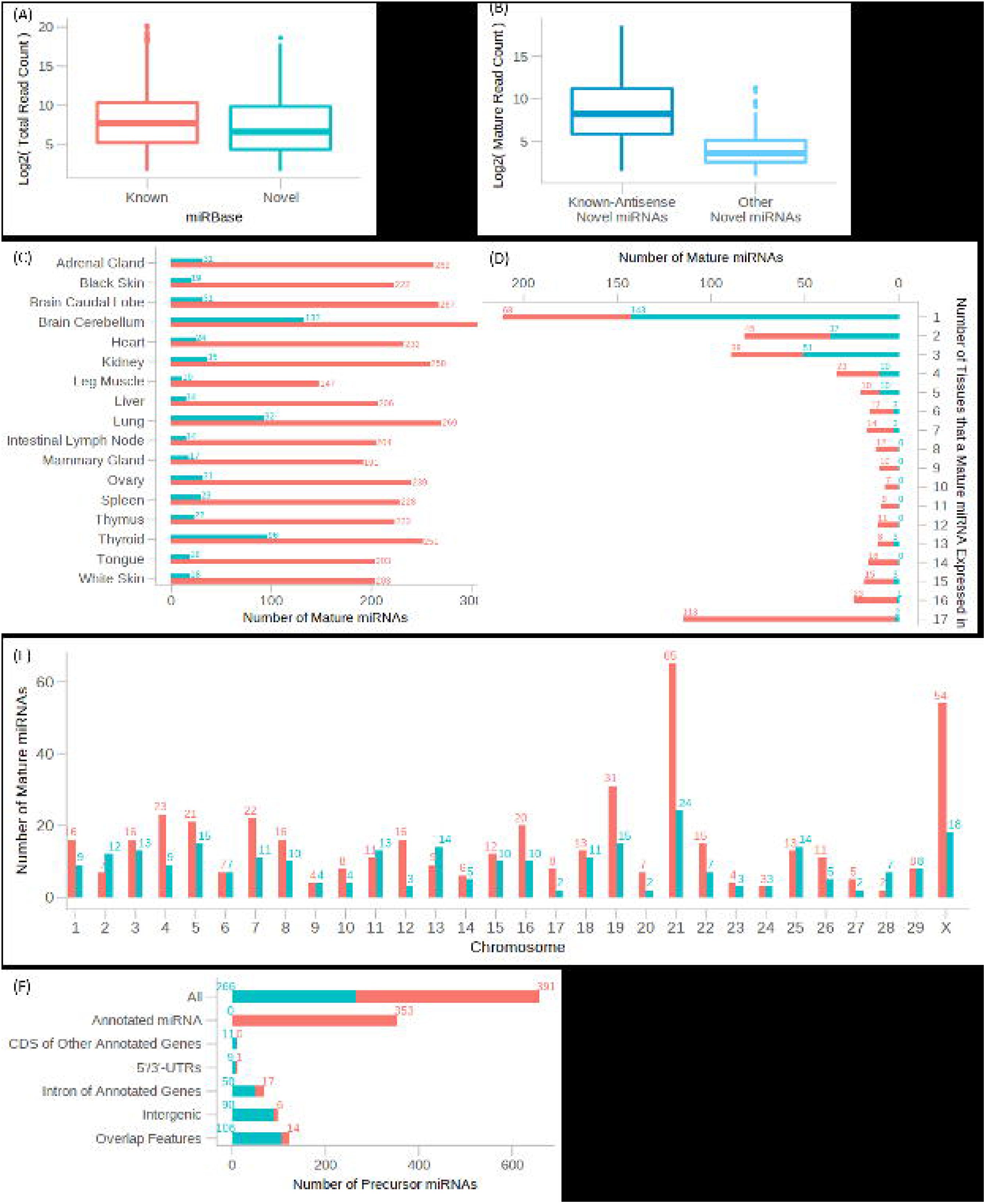
Summary of Expressed Known and Novel miRNAs. Red: known miRNAs. Blue: novel miRNAs. (A) Total read counts of mature miRNAs. (B) Read counts of ‘known-antisense’ novel mature miRNAs and other novel mature miRNAs. ‘Known-antisense’ novel miRNAs are novel miRNAs that overlapped with the antisense strand of known mature miRNA. (C) Number of mature miRNAs expressed in each tissue. (D) Number of tissues a mature miRNA was expressed in. (E) Number of mature miRNAs identified on chromosome 1-X. (F) Number of precursor miRNAs identified within genomic features annotated in UMD3.1.1. Order of annotation is shown in y-axis, i.e. annotated miRNAs, CDS, 5’-/3’-UTRs, introns, intergenic and overlapping features. Once a miRNA was assigned to a feature, the same miRNA was not counted again in the following features.

**Figure 2.**
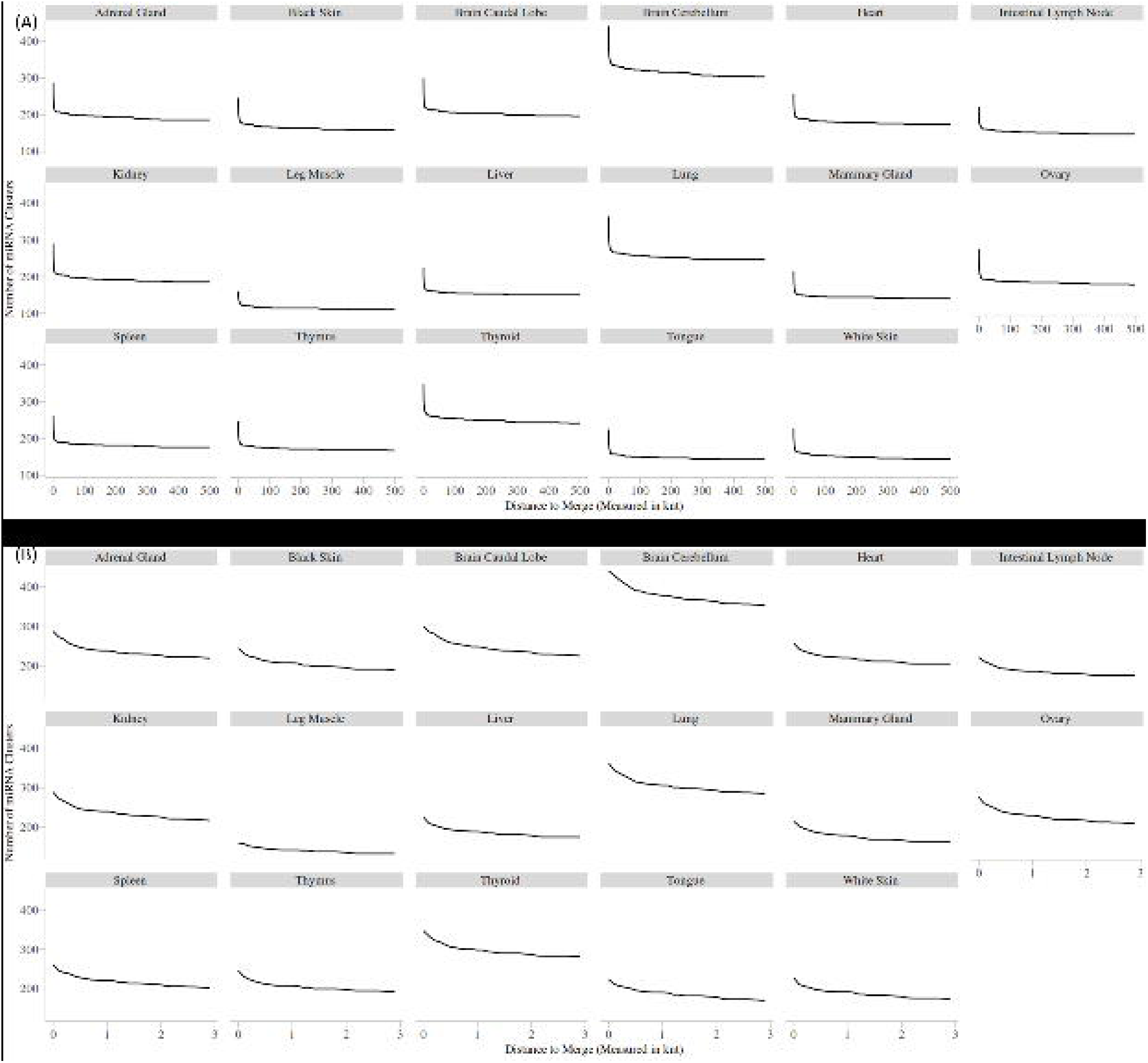
miRNAs Were Expressed in Clusters. Expressed miRNAs that are within a distance (x-axis) on the same DNA strand are merged into a “miRNA cluster”. Line plots shows the number of miRNA clusters (y-axis) in each tissue (title) within up to kilo-nucleotide (knt) distance, where distance ranges from Ont to 500knt (A) and 0nt to 3knt (B).

Close to 27% of the expressed mature miRNAs, which were known to miRBase (version 22) and were detected in our datasets, were not included in the General Feature Format (GFF) of the UMD3.1.1 reference genome annotation (e.g. *miR-654-5p, miR-6715-3p*). A small proportion of the expressed mature miRNAs (<1.5%), which had nearly identical sequences to annotated miRNAs in the GFF and miRBase (one or two nucleotides differences), were also identified, and those miRNAs were expressed from genomic regions that were not annotated in the GFF file. For example, *bta-miR-29d-3p (UAGCACCAUUUGAAAUCGAUUA)* is annotated on *chr16:77562591-77562612(+)* in GFF and this sequence is identical to that annotated in miRBase (version 22). In our dataset, 13 tissues (adrenal gland, black skin, brain caudal lobe, brain cerebellum, heart, kidney, lung, intestinal lymph node, mammary, ovary, thyroid, tongue and white skin) that had the mature miRNA *(uagcaccauuugaaaucgguua)* expressed were all from *chr16:77478646-77478667(+)*. Because *bta-miR-29d-3p* and *uagcaccauuugaaaucgguua* only differed in the 19^th^ nucleotide, miRDeep2 annotated *uagcaccauuugaaaucgguua* as *miR-29d-3p*. Overall, apart from the miRNAs that were already annotated in UMD3.1.1, expressed known and novel miRNAs were also found within the coding regions (CDS) of other annotated genes, 5’- or 3’-untranslated regions (UTRs), introns of annotated genes or non-coding RNAs, intergenic regions, and in genomic regions that crossed over from one of these features to another feature (Figure 1F), consistent with previous observations that miRNAs could arise throughout the genome (Melamed et al. 2013; Londin et al. 2015).

We traced the strand of the precursor that each mature and star miRNA were derived from, following the call of the miRNA community to provide a more precise nomenclature (Kozomara and Griffiths-Jones 2014; Budak et al. 2016) (Supplemental Table S2). We found that 83% of precursors had only one mature miRNA. We also observed some miRNAs had both arms highly co-expressed (read counts > 1000) in a tissue in both technical replicates, although most of those co-expressed duplexes were in lower abundance (read counts < 100). For example, *bta-mir-140* and *bta-mir-145* had both arms dominant in kidney (read counts: 1,088 to 13,635). *Bta-mir-361* had both arms dominant in adrenal gland, brain caudal lobe, brain cerebellum, heart, kidney, liver, ovary, spleen, thyroid and tongue (read counts: 223 to 1704). Some miRNAs changed the dominant arms across tissues. For example, *bta-miR-195-5p* was preferred in both technical replicates of black skin, lung, ovary, spleen and white skin (read counts: 230 to 3235), whereas *bta-miR-195-3p* was preferred in leg muscle, mammary gland and thyroid (read counts: 30 to 190). *Bta-miR-335-5p* was preferred in both technical replicates of thyroid (read count: 349 to 379), whereas *bta-miR-335-3p* was preferred in brain caudal lobe, heart, kidney, lung, ovary, spleen, tongue and white skin (read count: 24 to 464).

### Identification of differentially expressed mature miRNAs

Differential expression analysis was performed to identify mature miRNAs that were more often expressed (up-regulated) or less often expressed (down-regulated) in a tissue than the mean expression across all other tissues (Supplemental Table S3). Significant differentially expressed (after corrected for multiple testing P-value <0.01 and absolute fold change >2) mature miRNAs were observed in all tissues except in black skin (Table 2; Figure 3A). Known miRNAs with high tissue-specificity showed significant differential expression in our analysis, such as *miR-375* in adrenal gland (Ludwig et al. 2016; Gai et al. 2017), *miR-219* in brain caudal lobe and brain cerebellum (Ludwig et al. 2016), and *miR-122* in liver (Jopling 2012; Szabo and Bala 2013; Modepalli et al. 2014; Wang et al. 2016a). No novel miRNAs were found to be up-regulated in any tissues, but some novel miRNAs were down-regulated in adrenal gland, brain cerebellum, lung, ovary, spleen and thyroid. Tissues with similar biological function were grouped into clusters based on the full normalised read count matrix for all miRNAs (Figure 3B). The clustering profile of miRNAs was similar to that of mRNAs from the same tissues of the same cow shown previously (Chamberlain et al. 2015). For example, brain tissues and skin tissues were clustered together, respectively. However, some tissues were no longer grouped into the same cluster as they were in the mRNA profile. For example, leg muscle was separated from the muscle group formed by tongue and heart.

**Table 2.** Differentially Expressed miRNAs Observed in Each Tissue. Differentially expressed miRNAs are defined by P-value <0.01 and an absolute fold change >2 after correction for multiple testing. Novel miRNAs are represented by genomic coordinates. In tissues where no significant up- or down-regulated miRNAs was identified, they were not presented in the table.

| Tissue | Regulation | miRNAs of Significant Differential Expression |
| --- | --- | --- |
| Adrenal Gland | Up | *bta-miR-146b-5p, bta-miR-375-3p, bta-miR-409a-3p, bta-miR-543-3p, bta-miR-7-5p* |
|  | down | *bta-let-7c-5p, bta-miR-100-5p, bta-miR-181a-3p, bta-miR-183-5p, bta-miR-206-3p, bta-miR-215-5p, bta-miR-30c-5p, bta-miR-32-5p, bta-miR-340-3p, bta-miR-340-5p, bta-miR-532-5p, bta-miR-652-3p, bta-miR-98-3p, bta-miR-98-5p, chr8:83009655-83009675(-), chrX:96382704-96382725(+)* |
| Brain Caudal Lobe | Up | *bta-miR-212-5p, bta-miR-219-3p, bta-miR-323-3p* |
| Brain Cerebellum | up | *bta-miR-135b-5p, bta-miR-219-3p, bta-miR-219b-5p, bta-miR-323-3p* |
|  | down | *bta-miR-10a-5p, bta-miR-10b-5p, bta-miR-199b-5p, bta-miR-505-3p, bta-miR-505-5p, chr9:10768323-10768346(+)* |
| Heart | Up | *bta-miR-499-5p* |
| Intestinal Lymph Node | down | *bta-miR-361-5p* |
| Kidney | down | *bta-miR-101-3p, bta-miR-139-5p, bta-miR-150-5p, bta-miR-339b-5p, bta-miR-423-3p, bta-miR-92a-3p* |
| Leg Muscle | Up | *bta-miR-206-3p, bta-miR-378-3p, bta-miR-486-5p* |
| Liver | Up | *bta-miR-122-3p, bta-miR-122-5p, bta-miR-192-5p, bta-miR-194-5p* |
|  | down | *bta-miR-30c-5p* |
| Lung | Up | *bta-miR-34b-3p, bta-miR-34c-5p* |
|  | down | *chr1:79250542-79250563(+)* |
| Mammary Gland | down | *bta-miR-1271-5p* |
| Ovary | down | *chrX:34664642-34664663(+)* |
| Spleen | down | *chr8:83009655-83009675(-), chr9:10768323-10768346(+)* |
| Thymus | Up | *bta-miR-106a-5p* |
|  | down | *bta-miR-193b-3p, bta-miR-671-5p, bta-miR-193b-5p, bta-miR-493-5p* |
| Thyroid | Up | *bta-miR-135b-5p* |
|  | down | *chr18:58014874-58014895(-), chr7:39194962-39194983(-)* |
| Tongue | Up | *bta-miR-206-3p* |
| White Skin | down | *bta-let-7b-3p* |

**Figure 3.**
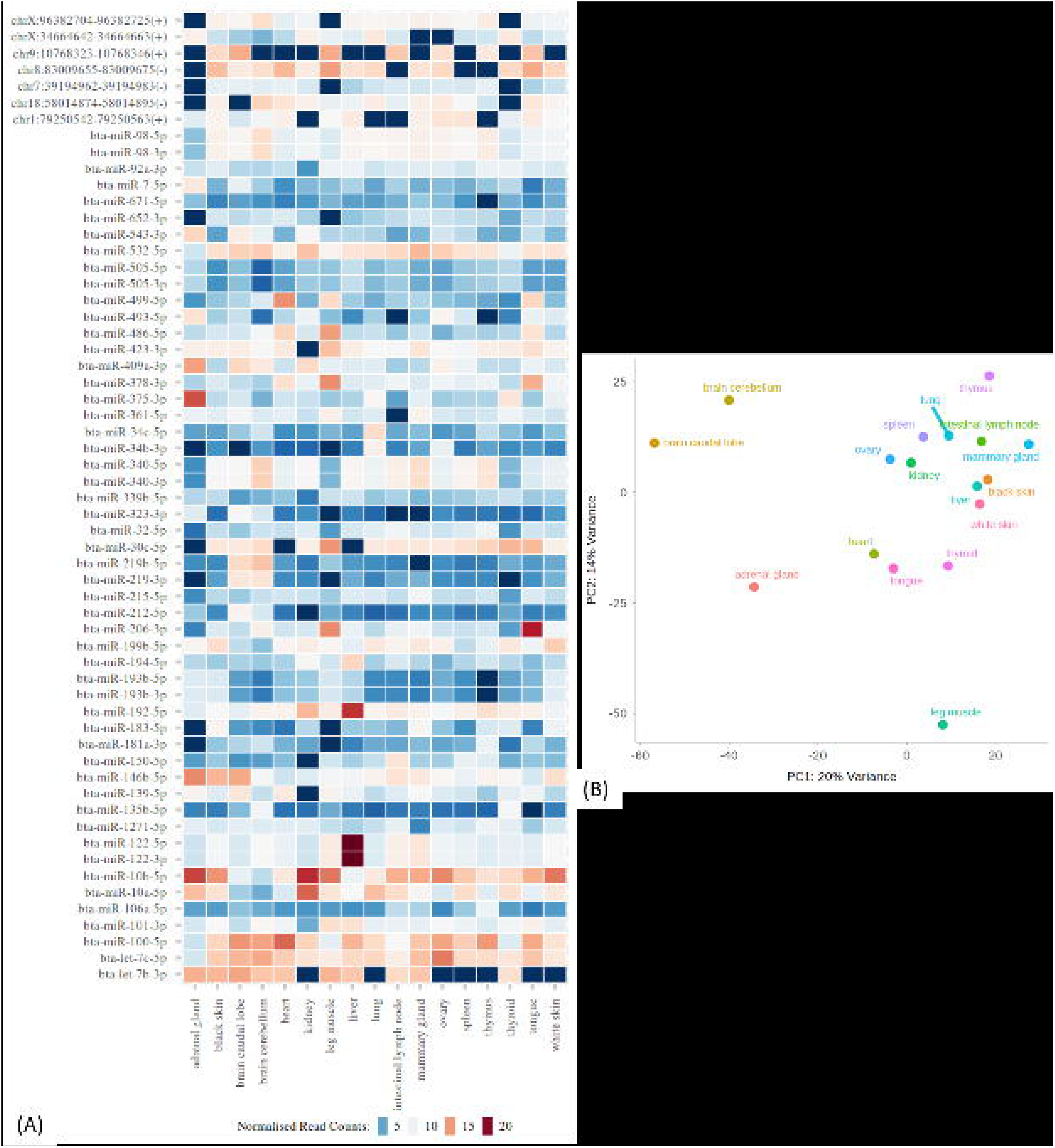
Mature miRNA Expression Patterns. (A) A heatmap showing mature miRNAs (y-axis) that were significantly differentially expressed (after correction for multiple testing P-value < 0.01 and an absolute fold change > 2) in a bovine tissue (x-axis) compared with all other bovine tissues. Colour key indicates normalised read counts of the mature miRNA detected in a tissue by miRDeep2. Red: Up-regulated; Blue: Down-regulated. (B) A principle component analysis (PCA) plot showing tissue- to-tissue distance measured by expression of all mature miRNAs.

### Prediction of miRNA seed match sites

A miRNA seed match site is a segment of mRNA sequence that perfectly reverse-complements the miRNA seed sequence, which is the 2^nd^ to 7^th^ nucleotide at the 5’-end of the mature miRNA sequence. To identify miRNA seed match sites within bovine mRNA transcripts for our expressed mature miRNAs, all clean (Materials and Methods) mature miRNA sequences were provided as input to TargetScan (Agarwal et al. 2015) (version 7.2). This consisted of 434 known and 265 novel mature miRNA sequences, which were expressed in at least one of the 17 bovine tissues. Based on the identical extended seed sequence (i.e. 2^nd^-8^th^ nucleotides at the 5’-end of mature miRNA sequence), all mature miRNAs were grouped into 600 miRNA families. Examining members within each miRNA family, we found that 455 families were formed by grouping known and novel expressed miRNAs with miRNAs in miRBase (Kozomara and Griffiths-Jones 2014) (version 22), 125 families were formed by a single expressed novel miRNA that was not grouped with any miRBase miRNAs, and the remaining 20 families were formed by a single known expressed miRNA that was also not grouped with any miRBase miRNAs. Most of these “ungrouped” known miRNAs were recently discovered, as the name of these miRNAs had large numbers (e.g. *bta-miR-11995-3p*) as opposed to small numbers (e.g. *bta-miR-1*), noting miRNAs are mostly named in chronological order of their discovery (Kozomara and Griffiths-Jones 2014).

TargetScan (Agarwal et al. 2015) (version7.2) returned information including the genomic position of each seed match site of a miRNA family in bovine reference genome bostau7 that was orthologous to the 3’UTR of mRNA transcripts in human reference genome (UCSC ID) hg19. Although TargetScan only predicted miRNA seed match sites in the 3’UTRs, we searched through all genes in bostau6 for seed match sites, because miRNAs were found to be reverse-complements to the seed match sequences in the promoter sequence, 5’UTR and open reading frames (Lewis et al. 2005; Stark et al. 2007; Place et al. 2008; Schnall-Levin et al. 2010; Cheng et al. 2015; Xiao et al. 2016).

Our prediction using TargetScan returned a total of 6,411,460 records of “mRNA target: seed match site: miRNA family” from bostau7. Among those, 879,240 records were recovered (Materials and Methods) on the 31 standard chromosomes (1-29, X and M) from bostau6. This included a total of 780,481 putative seed match sites from 18,196 mRNA transcripts from 16,381 genes on the 31 chromosomes. Although TargetScan returned putative seed match sites on bostau7 that was at least 6-8nt long, because only the seed match sites were searched on bostau6, each putative seed match site on bostau6 was 6nt. A miRNA family was predicted to target 4 to 18,448 sites (on average 1,306) from 4 to 7,591 mRNA transcripts (on average 1,292) from 4 to 6,767 genes (on average 1,154). A target gene was predicted to have 1 to 199 miRNA seed match sites (on average 48). Given the many-to-many relationship among miRNA family and mRNA, a total of 692,685 putative interactions between miRNA families and cognate gene targets were returned from our prediction on bostau6, which resulted in 3,484,895 putative interactions between expressed miRNAs and cognate gene targets. Genomic coordinates of all predicted seed match sites of our expressed miRNAs are reported (Supplemental Table S4).

Acetylated lysine 27 on histone H3 (H3K27ac) and tri-methylation of lysine 4 on histone H3 (H3K4me3) regions have been identified by chromatin immunoprecipitation followed by high-throughput sequencing (ChIP-Seq) experiments to identify active enhancer regions and active promoter regions (Creyghton et al. 2010; spicuglia and Vanhille 2012). H3K27ac and H3K4me3 ChlP-Seq marks from bovine liver tissue were mostly found at intergenic and intronic regions, and were also found at 5’UTRs, 3’UTRs and CDS (Wang et al. 2017). In the nucleus, miRNAs were shown to bind to the promoter sequence through the same 6nt miRNA seed match site to increase transcription of the gene that the promoter regulates (Place et al. 2008; Zhang et al. 2014; Cheng et al. 2015; Xiao et al. 2016). This inspired us to examine within genes, whether DNA sequences that were marked by bovine H3K27ac or H3K4me3 signals (Villar et al. 2015; Zhao et al. 2015) were more or less often also putative miRNA seed match sites.

To answer this question, a contingency table was constructed to show the number of nucleotides within gene regions in UMD3.1 (Ensembl release 91) that were (1) inside both a histone modification mark and putative miRNA target sites, (2) inside a histone modification mark but outside putative miRNA target sites, (3) outside a histone modification mark but inside putative miRNA target sites, and (4) outside both a histone medication mark and putative miRNA target sites (Table 3). Because putative miRNA seed match sites were DNA-strand specific, where a histone modification mark existed we assumed both strands affected. Using data from the contingency table, a Chi-Square test showed that within genes, the proportion of miRNA seed match sites that were also within the histone modification mark regions was significantly higher than the proportion of non-miRNA seed match sites that were within the same histone mark regions (P-value <0.00001; Table 3), indicating that miRNA seed match sites preferentially occurred within gene regions undergoing active histone modifications.

**Table 3.** Enrichment of miRNA Seed Match Sites within Bovine-specific Histone Modification Signals. The number of nucleotides inside/outside all putative miRNA seed match sites and inside/outside a histone modification signal. Also, the proportion of nucleotides inside/outside miRNA seed match site that were inside a histone modification signal showing the direction of the test, and the Chi-Square test results showing the significance of the test.

| H3K27ac (Liver) (Villar et al. 2015) |  |  |  |  |  |
| --- | --- | --- | --- | --- | --- |
|  |  | Inside | Outside | Total | Prop. Inside |
| miRNA Seed Match Sites | Inside | 532400 | 3811824 | 4344224 | 12.2554% |
|  | Outside | 40646950 | 674559111 | 715206061 | 5.6833% |
|  | Total | 41179350 | 678370935 | 719550285 |  |
| Chi-Square Test | $\chi^2$-value | | | 345673.9 | |
|  | P-value |  |  | 0 |  |

| H3K4me3 (Liver) (Villar et al. 2015) |  |  |  |  |  |
| --- | --- | --- | --- | --- | --- |
|  |  | Inside | Outside | Total | Prop. Inside |
| miRNA Seed Match Sites | Inside | 343365 | 4000859 | 4344224 | 7.9039% |
|  | Outside | 14275698 | 700930363 | 715206061 | 1.9960% |
|  | Total | 14619063 | 704931222 | 719550285 |  |
| Chi-Square Test | $\chi^2$-value | | | 757193.9 | |
|  | P-value |  |  | 0 |  |
| H3K4me3 (Tender Muscle) (Zhao et al. 2015) |  |  |  |  |  |
|  |  | Inside | Outside | Total | Prop. Inside |
| miRNA Seed Match Sites | Inside | 75891 | 4268333 | 4344224 | 1.7469% |
|  | Outside | 9127502 | 706078559 | 715206061 | 1.2762% |
|  | Total | 9203393 | 710346892 | 719550285 |  |
| Chi-Square Test | $\chi^2$-value | | | 7577.71 | |
|  | P-value |  |  | 0 |  |
| H3K4me3 (Tough Muscle) (Zhao et al. 2015) |  |  |  |  |  |
|  |  | Inside | Outside | Total | Prop. Inside |
| miRNA Seed Match Sites | Inside | 89620 | 4254604 | 4344224 | 2.0630% |
|  | Outside | 10549060 | 704657001 | 715206061 | 1.4750% |
|  | Total | 10638680 | 708911605 | 719550285 |  |
| Chi-Square Test | $\chi^2$-value | | | 10248.99 | |
|  | P-value |  |  | 0 |  |

### Confirmation of putative interactions between miRNAs and targets

Putative interactions between miRNA families and cognate targets were confirmed by interactions identified from previous publications through experimental and computational procedures (Table 4). Messenger RNA (mRNA) sequences bound with miRNAs by the AGO protein in bovine kidney cells were identified through CLIP-Seq (Scheel et al. 2017). After converting genomic coordinates of those mRNA sequences from (UCSC ID) bostau7 to bostau6, 208,688 interactions from 224 expressed miRNAs and 200,459 cognate target sequences remained. We identified 297,802 putative interactions from 238 miRNA families and 15,999 seed match sites in our lactating cow’s kidney tissue. Of those, 11,885 putative interactions from 163 miRNA families and 9,282 seed match sites overlapped with those from the Scheel *et al*. set (Table 4). Other miRNA interaction datasets from public domains did not provide any cell/tissue information, and therefore all our putative interactions were counted and compared. We found 3,173 putative interactions between miRNA families and target gene names overlapped with that in miRTarBase (Chou et al. 2018), 229,473 putative interactions between miRNA families and target mRNA transcript IDs overlapped with that in miRWalk (Dweep and Gretz 2015), and 182,685 putative interactions between miRNA families and target gene names overlapped with that in TargetScan (Agarwal et al. 2015) (version 7) (Table 4).

**Table 4.** Number of Confirmed Putative Interactions Between miRNA Family and Cognate Targets. Presented are the number of interactions between miRNAs and cogante target sequences, target gene names or target transcript IDs in confirmation databases. Also presented are the number of interactions from miRNA families and cognate seed match sites, target gene names or target transcript IDs in our prediction. Last presented are the number of interactions that overlapped between the confirmation set and our prediction set. Genomic coordinates of miRNA target sequences (Scheel et al. 2017) were converted from bostau7 to bostau6 prior to counting and only interactions in kidney were counted. Mammalian target gene names in miRTarBase (Chou et al. 2018) and TargetScan (Agarwal et al. 2015) were converted to bovine orthologs and all NA records were removed prior to counting. Apart from CLIP-Seq (Scheel et al. 2017), no tissue information is provided in any other public datasets, and therefore all interactions were counted.

| Type | # miRNA | # Target | # Interactions |
| --- | --- | --- | --- |
| CLIP-Seq (Scheel et al. 2017) | 224 miRNAs | 200459 sequences | 208688 |
| Our Prediction | 238 miRNA families | 15999 sites | 297802 |
| Overlaps | 163 miRNA families | 9282 sites | 11885 |
| miRTarBase (Chou et al. 2018) | 950 miRNAs | 8242 gene names | 20538 |
| Our Prediction | 600 miRNA families | 14947 gene names | 651674 |
| Overlaps | 347 miRNA families | 7706 gene names | 3173 |
| miRWalk (Dweep and Gretz 2015) | 794 miRNAs | 17022 transcript IDs | 6838356 |
| Our Prediction | 600 miRNA families | 18196 transcript IDs | 775378 |
| Overlaps | 358 miRNA families | 14909 transcript IDs | 229473 |
| TargetScan (Agarwal et al. 2015) | 2407 miRNAs | 15532 gene names | 7440300 |
| Our Prediction | 600 miRNA families | 14947 gene names | 651674 |
| Overlaps | 319 miRNA families | 14908 gene names | 182685 |

We investigated the relationship between miRNA transcription and target mRNA transcription. mRNA transcripts that were differentially expressed in a tissue compared with the mean expression in all other tissues (after corrected for multiple testing P-value <0.01 and absolute fold change >2) from the same cow as used in this study and previously identified (Chamberlain et al. 2015) were utilised. Combining those results with differentially expressed miRNAs that were identified in this study, we found that 84,585 tissue-specific putative interactions between miRNA families and target genes had both a miRNA in the miRNA family and the gene target differentially expressed in the same tissue from the same cow. Of those 84,585 interactions, which consisted of 15 tissues, 2,333 target gene IDs and 473 miRNA families, 233 putative interactions from all tissues that were confirmed by miRTarBase (Chou et al. 2018) and 8 putative interactions in kidney that were confirmed by CLIP-Seq (Scheel et al. 2017) had both the miRNA and gene differentially expressed in the same tissue, and one putative interaction overlapped the miRTarBase, CLIP-Seq and differential expression sets. KEGG and GO analyses on differentially expressed mRNAs were previously performed (Chamberlain et al. 2015). Combining functional terms from KEGG and GO, we found that mRNAs that were co-differentially expressed with cognate miRNA within the same tissue were most-frequently related to metabolism or metabolic processes, immune or inflammatory responses, signalling pathways, cell activation and differentiations, blood coagulation and transportations of inorganic and organic molecules (Chamberlain et al. 2015).

To investigate whether there was a general direction between miRNA transcription and target mRNA transcription, a least square analysis was performed to test the association between the Iog2 fold change of miRNA transcription from this study and the log2 fold change of mRNA transcription from Chamberlain *et al*. for miRNA and target mRNAs that were both significantly differentially-expressed (after correction for multiple testing P-value <0.01 and absolute fold change >2) in the same tissue of the same cow. The number of records for the fold change of expression between miRNAs and target mRNAs across tissues varied from 62 to 95,586. We found no significant (P-value < 10^−s^) association between the fold change of miRNA transcription and target mRNA transcription in any tissues (Table 5). Note that although leg muscle and lung appeared to be significant, manual examination of the data did not support a true correlation. These results indicated that in general if miRNAs were up-regulated in a tissue, their target mRNAs could be up- or down-regulated in the same tissue.

**Table 5.** Correlation Between miRNAs and mRNAs of Significant Differential Expression in the same tissue. The correlation between the Log2 fold change of mRNA expression (Chamberlain et al. 2015) and the Log2 fold change of miRNA expression in this study within each tissue from a single dairy lactating cow was tested. The coefficient and p-value from correlation test within each tissue are presented. In some tissues, the coefficient and p-value were NA because only one differentially expressed (after corrected for multiple testing P-value <0.01 and absolute fold change >2) miRNA was found within a tissue.

| Tissue | Effect | P-value |
| --- | --- | --- |
| Adrenal Gland | 0.003091 | 0.392132 |
| Brain Caudal Lobe | 0.745447 | $9.69 \times 10^{-7}$ |
| Brain Cerebellum | -0.00281 | 0.072024 |
| Heart | NA | NA |
| Kidney | 0.053872 | 0.106306 |
| Leg Muscle | -4.59879 | $6.76 \times 10^{-9}$ |
| Liver | -0.07229 | $7.78 \times 10^{-5}$ |
| Lung | 0.05387 | $1.15 \times 10^{-9}$ |
| Mammary Gland | NA | NA |
| Ovary | NA | NA |
| Spleen | $-5.2 \times 10^{-17}$ | 1 |
| Thymus | -0.07477 | 0.001095 |
| Thyroid | -0.00552 | 0.492063 |
| Tongue | NA | NA |
| White Skin | NA | NA |

Mature miRNAs that were derived from different arms of the same precursor were found to target different genes (Supplemental Table S5). *Bta-miR-140-5p, bta-miR-140-3p, bta-miR-145-5p, bta-miR-145-3p, bta-miR-195-5p, bta-miR-195-3p, bta-miR-335-5p, bta-miR-335-3p, bta-miR-361-5p* and *bta-miR-361-3p* were respectively the dominant product in at least one tissue from this lactating cow (Supplemental Table S2). These mature miRNAs were predicted to target 1431, 1712, 770, 897, 5868, 521, 1437, 3415, 411 and 1932 bovine mRNA genes, respectively. Among those, 24, 16, 11, 3, 1432, 3, 192, 2, 11 and 10 target genes were confirmed by experimental procedures that were published in either miRTarBase (Chou et al. 2018) (all tissues were considered) or CLIP-Seq (Scheel et al. 2017) (only kidney was considered).

### Enrichment analysis

A permutation test was performed to examine whether putative interactions between the miRNA families and cognate targets were significantly enriched for interactions identified from experimental and computational procedures in previous publications (Table 6). We found that putative interactions were statistically significantly enriched for previously-identified interactions from all confirmation datasets. Compared with 10,000 random interactions, putative interactions that were confirmed by miRTarBase (Chou et al. 2018) were 1.61-fold higher, CLIP-Seq (Scheel et al. 2017) were 1.19-fold higher, miRWalk (Dweep and Gretz 2015) were 1.11-fold higher, TargetScan (Agarwal et al. 2015) were 1.40-fold higher, and by the differential expression of mRNAs (Chamberlain et al. 2015) and miRNAs from the same tissue of the same cow were 1.10-fold higher (Table 6).

**Table 6.** Enrichment of Confirmed Putative Interactions Between miRNA Families and Cognate Targets. Significance is denoted as “<0.0001” if the putative interactions are more often confirmed by interactions from a confirmation dataset than all 10,000 randomly-shuffled permutations; otherwise significance is denoted as the ranking of the actual degree of overlap among the list of 10,000 random overlaps. Fold change of enrichment is the ratio of the actual number of confirmed putative interactions to the average number of confirmed random interactions from 10,000 permutations.

| Confirmation Dataset | Significance | Fold Change |
| --- | --- | --- |
| miRTarBase (Chou et al. 2018) | <0.0001 | 1.607254 |
| CLIP-Seq (Scheel et al. 2017) | <0.0001 | 1.193042 |
| miRWalk (Dweep and Gretz 2015) | <0.0001 | 1.112938 |
| TargetScan 7.2 (Agarwal et al. 2015) | <0.0001 | 1.398332 |

### Polymorphisms within miRNA genes and targets

To investigate the possibility that polymorphism within miRNA genes and target sites might be detrimental for fitness and therefore could be a target for natural selection, we compared both the rate of polymorphism and the allele frequency distributions within miRNA genes and targets with that across the genome.

Firstly, we defined the polymorphic rate as the proportion of a genomic feature that were polymorphic sites. We used whole genome sequence (WGS) variants from the 1000 Bull Genomes Project (Daetwyler et al. 2014) (*Bos Taurus* Run 6) that were filtered based on quality metrics as described (Daetwyler et al. 2017). The filtered variants (including single-base-nucleotide substitutions, insertions and deletions) consisted of sequence variants with high confidence, which have been shown to improve the quality of variant sets judged by the concordance of sequence and SNP chip genotypes at overlapping positions as well as the rate of opposing homozygotes found in parent-offspring pairs (Daetwyler et al. 2017). We found that polymorphic rates were lower within miRNA genes (precursor, mature and star) and miRNA targets (target genes and seed match sites) than the polymorphic rate averaged across the genome (Table 7). For mRNA genes that were targeted by our expressed miRNAs, their gene regions (including introns) had a higher polymorphic rate than 3’UTRs. miRNA seed match sites from *in-silico* prediction and miRNA target sequences from previous experimental identification (Scheel et al. 2017) had even lower polymorphic rates than 3’UTRs. Compared with genes that were targeted by expressed miRNAs (1.556%), miRNA genes (0.710%) were much more depleted for polymorphic sites. Known expressed miRNAs had a lower polymorphic rate than novel expressed miRNAs. Mature miRNAs had the lowest polymorphic rate among all genomic features examined. Surprisingly, although star miRNAs were partially reverse-complementary with mature miRNAs, the star sequence had a higher polymorphic rate rate than the mature sequence (Table 7).

**Table 7.** Density of WGS Variants within Genomic Features. The density of filtered whole genome sequence variants as described by Daetwyler *et al*. (Daetwyler et al. 2017) was the proportion of a genomic feature being polymorphic (including both SNPs and INDELs). The variants were identified from the 1000 Bull Genomes Project (Daetwyler et al. 2014) Bos Taurus Run 6. mRNA Genes Targeted by Expressed miRNAs 1.556% Computationally Predicted miRNA Seed Match Sites 1.375% Table 8 Density of Raw WGS Variants with <1% Allele Frequency within Genomic Features. The density was the proportion of raw single-base-nucleotide substitution variants within a genomic feature with an allele frequency less than 1%. The raw whole genome sequence variants were identified from the 1000 Bull Genomes Project (Daetwyler et al. 2014) (*Bos Taurus* Run 6).

| Genomic Feature |  | Density |
| --- | --- | --- |
| Genome Wide |  | 1.679% |
| 3'UTRs Genome-Wide |  | 1.416% |
| Known Expressed miRNA Genes | Precursor | 0.710% |
|  | Mature | 0.619% |
|  | Star | 0.708% |
| Novel Expressed miRNA Genes | Precursor | 1.223% |
|  | Mature | 1.117% |
|  | Star | 1.225% |
| Experimentally Identified | miRNA Target Sequences (Scheel et al. 2017) | 1.376% |
| Computationally Predicted | mRNA Genes Targeted by Expressed miRNAs | 1.556% |
|  | miRNA Seed Match Sites | 1.375% |

All alleles in the variant call file, including each alternative allele at the same polymorphic site, were calculated for their frequencies in the 2,333 animals from the 1000 Bull Genomes Project (Daetwyler et al. 2014) (*Bos Taurus* Run 6). To examine whether miRNA genes and target sites had more variants with extreme frequencies than the entire genome, which could be evidence of selection at these genomic features, we used raw WGS variants (single-base-nucleotide substitutions only). Raw WGS variants contained all variants called, so rare variants were not filtered out. We found that variants with extreme allele frequencies were more likely within mature, star and precursor miRNAs, followed by miRNA target sequences from experimental identification (Scheel et al. 2017) and putative miRNA seed match sites within target genes from computational prediction, compared to the entire genome (Table 8; Figure 4).

**Table 8.**
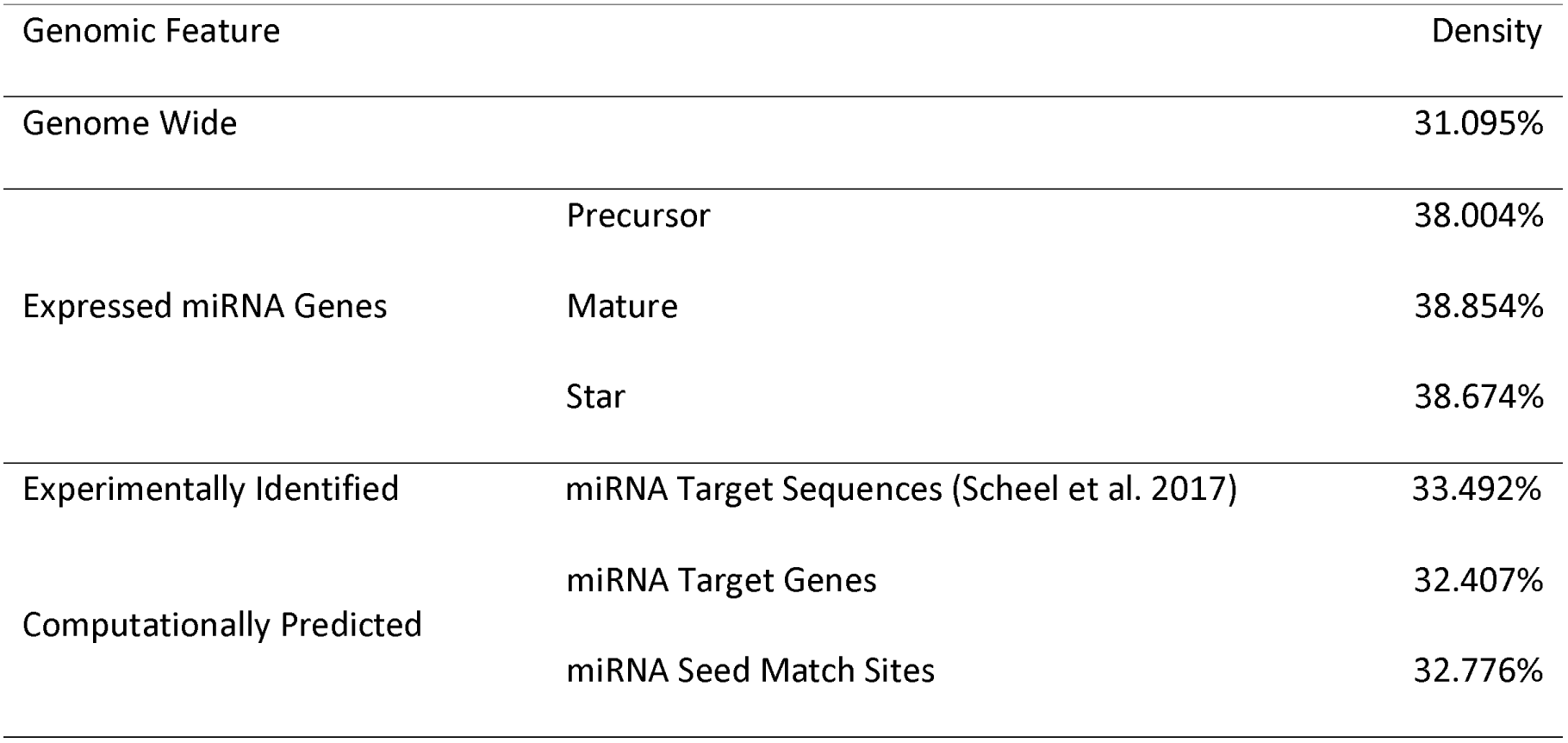
Density of Raw WGS Variants with <1% Allele Frequency within Genomic Features. The density was the proportion of raw single-base-nucleotide substitution variants within a genomic feature with an allele frequency less than 1%. The raw whole genome sequence variants were 1097 identified from the 1000 Bull Genomes Project (Daetwyler et al. 2014) (*Bos Taurus* Run 6).

| Genomic Feature |  | Density |
| --- | --- | --- |
| Genome Wide |  | 31.095% |
| Expressed miRNA Genes | Precursor | 38.004% |
|  | Mature | 38.854% |
|  | Star | 38.674% |
| Experimentally Identified | miRNA Target Sequences (Scheel et al. 2017) | 33.492% |
| Computationally Predicted | miRNA Target Genes | 32.407% |
|  | miRNA Seed Match Sites | 32.776% |

**Figure 4.**
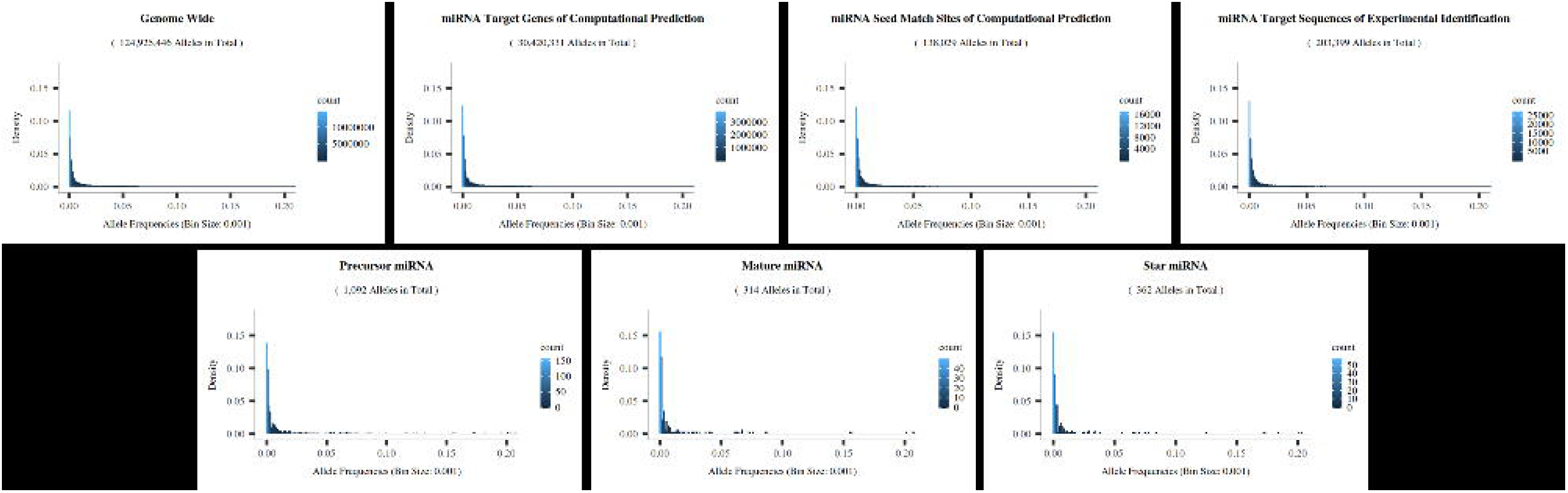
Allele Frequencies for Raw WGS Variants within miRNA Genes, miRNA Targets and the Entire Genome. Histogram shows the proportion of alleles (y-axis) within a genomic feature (title) that fell within a frequency range (x-axis). The height of each bar was the number of alleles (legend) within a frequency range divided by the total number of alleles within the genomic feature (subtitle). For each allele present among the 2,333 animals from 1000 Bull Genomes Project (Daetwyler et al. 2014) (*Bos Taurus* Run 6), including different alleles at the same polymorphic site, its frequency was calculated, but only allele frequencies less than 20% from single-base-nucleotide substitution variants were plotted.

Insertions and deletions (INDELs) were categorised by the number of positional shifts, *n ∊* (−∞, +∞), between the alternative allele and reference allele at the same locus. To examine whether INDELs with any n-th positional shift were more enriched within miRNA genes and target sites than the entire genome, we used the raw INDELs (unfiltered) from 1000 Bull Genomes Project (Daetwyler et al. 2014) (*Bos Taurus* Run 6). We found that most INDELs only shifted a small number of positions (Figure 5), which was consistent with previous research in humans that INDELs with a small number of positional shifts were more frequent than INDELs with a large number of positional shifts, even though those studies did not use WGS variants (Mills et al. 2006; Saunders et al. 2007; Bhattacharya et al. 2012; Gong et al. 2012). Compared with the entire genome and mRNA genes that were targeted by our expressed miRNAs, miRNA genes were highly depleted for INDELs of all lengths. Interestingly, both miRNA seed match sites from our computational prediction and miRNA target sequences from experimental identification (Scheel et al. 2017) had a higher density of INDELs than the entire genome and target mRNA genes (Figure 5). INDELs with a single base pair positional shift and even number of positional shifts were more dominant in putative miRNA seed match sites. Deletions were more frequently observed in experimentally identified miRNA target sequences of Scheel *et al*. (2017) than insertions.

**Figure 5.**
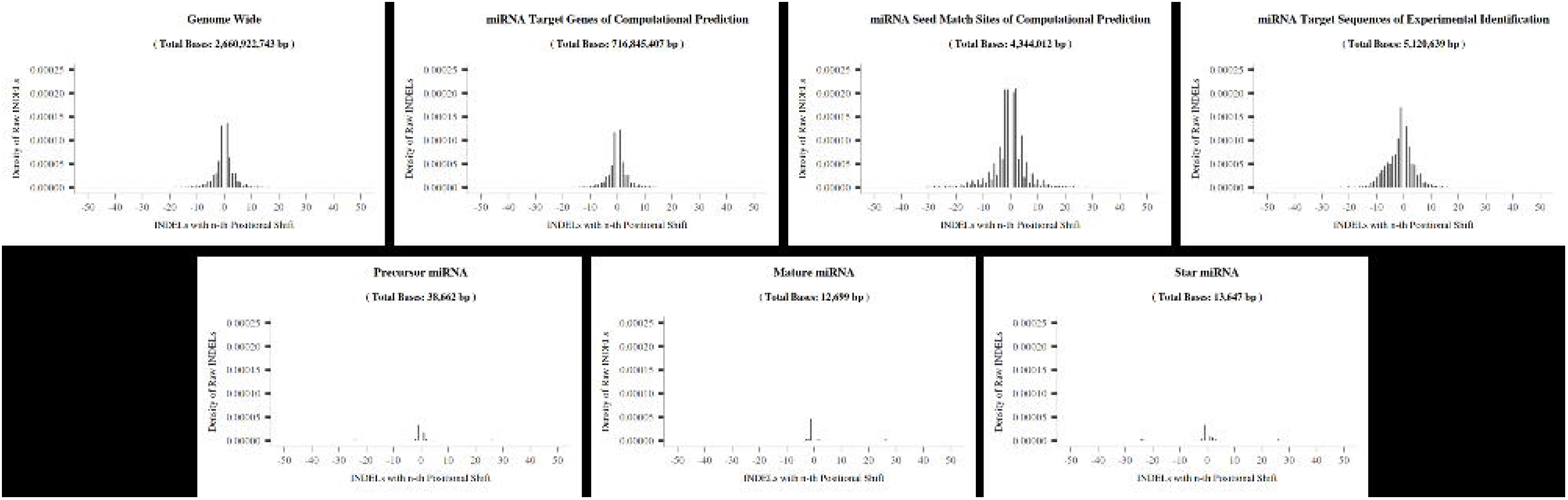
Frequency of an *n*-th Positional Shift within miRNA Genes, miRNA Targets and the Entire Genome. Raw sequence variants (INDELs only) were identified from whole genome sequencing of 2,333 key ancestor *Bos Taurus* bulls from 1000 Bull Genomes Project (Daetwyler et al. 2014) (Run 6). Each bar (y-axis) represented the number of INDELs with an ***n***-th positional shift (x-axis) divided by the size (subtitle) of the genomic feature (title). An ***n***-th positional shift is the difference between the number of nucleotides in the alternative allele to the number of nucleotides in the reference allele at the same locus. If >1 alternative allele at a locus was detected, each alternative allele was calculated. Only ***n ∊*(−50, 0)** & **(0, 50)** were shown. The genomic coordinates of experimentally identified miRNA target sequences (Scheel et al. 2017) were converted from bostau7 to bostau6 before calculation.

### Examination of associations among miRNAs, miRNA target regions and allele-specific expression of target genes

Allele-specific expression (ASE) or allelic imbalance is the biased expression from one parental allele for a gene. In 18 tissues from this cow, the degree of allelic imbalance at a locus was previously reported as a Chi-Square value (Chamberlain et al. 2015). To examine whether allelic imbalance was associated with the miRNAs and miRNA target sites, we defined ASE score as the square root of this Chi-square value, and putative miRNA target sequence as the 56nt mRNA sequence surrounding the miRNA seed match site (Materials and Methods).

Firstly, we examined across tissues, whether allelic imbalance at each heterozygous site within putative miRNA target sequence was associated with the expression level of cognate mature miRNA. A dataset with 176,478 records was constructed, which consisted of 682 mature miRNAs and 17,626 heterozygous sites within 19,119 cognate putative target sequences in 6,704 target genes across 17 tissues. Results of analysis of variance (ANOVA) showed that the expression level of miRNA did not have a significant effect on the allelic imbalance within its putative miRNA target sequence (P-value = 0.367).

Secondly, we examined across genes, whether zygosity within putative miRNA target sequences was associated with allelic imbalance within exons of miRNA target genes. A dataset with 8,406,848 records was constructed, which consisted of 191,866 WGS variants from 1000 Bull Genomes Project (Daetwyler et al. 2014) (*Bos Taurus* Run 5) within 158,466 putative miRNA target sequences from 7,072 miRNA gene targets, and ASE scores from 22,238 heterozygous sites within exons of miRNA target genes. Results from linear regression showed that zygosity within putative miRNA target sequences had an effect on allelic imbalance within exons of target genes. Also, with an addition of a heterozygous site within a putative miRNA target sequence, we could expect the mean allelic imbalance within exons of miRNA target genes to increase 0.0389 (P-value ≤ 10^−8^).

A permutation test was performed to assess whether the effect of zygosity within putative miRNA target sequences was significant compared with the effect of zygosities within genes that were not putative miRNA target sequence nor exons (termed the ‘null regions’). Results showed that all effects from the null regions were positive. Additionally, the effect of zygosity within putative miRNA target sequences, 0.038937, was only larger than 845 out of 10,000 effects from the null regions. The observed standard error of the effect within putative miRNA target sequences, 0.002797, was bigger than 9,998 out of 10,000 standard errors from null regions. This showed that an increment of polymorphic sites anywhere within a gene increased allelic imbalance in exons, and zygosity within putative miRNA target sequences were not more significantly associated with allelic imbalance at exons of target genes than zygosity within the gene but outside putative miRNA target sequence and exons.

## Discussion

Improving the annotation of functional elements (or regulatory elements) in the bovine genome will help to better understand the effects of sequence variants across the genome, to fill the gap between genotype and phenotype, and to contribute to the application of molecular phenotype to predict complex phenotypes (Andersson et al. 2015). A functional element is denoted as a discrete genomic segment that encodes a defined product (e.g. protein, mRNA or miRNA) or a reproducible biochemical signature (e.g. protein binding sites or miRNA seed match sites) (ENCODE Project Consortium 2012). In this study, we improve the annotation of tissue-specific miRNAs that were expressed in a single lactating dairy cow. We provide a list of putative seed match sites for all expressed mature miRNAs and propose possible functions of those miRNA-led interactions.

Most of our findings on expressed miRNAs aligned with previous studies in human and other animals. First, we confirmed the expression of many widely-reported, tissue-specific and highly-expressed miRNAs (Supplemental Table S2), such as myotubes-related *miR-1* and *miR-133b* (Lim et al. 2005; Zhao et al. 2005) in heart, leg muscle and tongue as well as neuron-specific *miR-9* (Wang et al. 2012) in brain caudal lobe and brain cerebellum. Second, a large proportion of miRNAs identified in this study were novel, and overall had a lower expression level but a higher tissue-specificity than known miRNAs (Figure 1) (Friedländer et al. 2012; Londin et al. 2015; Huang et al. 2017). Novel miRNAs tended to be expressed from the antisense strand of known miRNAs (Figure 1), which were suggested to have functional importance in different stages of development (Bender 2008; Stark et al. 2008; Tyler et al. 2008). Finally, most expressed miRNAs were close to one another on the DNA strand, forming into discrete “blocks” or “clusters” (Figure 2). miRNA clusters were previously defined as a set of two or more miRNAs that are transcribed in the same orientation and are not separated by a transcription unit or a miRNA in the opposite orientation (Altuvia et al. 2005). miRNA clusters are known to play important roles controlling various cellular process in the human genome (Yu et al. 2006; Griffiths-Jones et al. 2008; Lai and Vera 2013; Marco et al. 2013). Constituents of the same miRNA cluster were suggested to be expressed from the same transcription unit and evolve to coordinate regulation of functionally related genes (Marco et al. 2013; Wang et al. 2016b). Overall, our results suggested that bovine miRNAs were more extensive than currently represented by public repositories, and we have characterised a significant number of novel tissue-specific miRNAs.

We observed cases of identical miRNA sequences being expressed at different magnitudes from different genomic locations in different tissues. This tissue-specificity of miRNA expression could be controlled by super-enhancers (SEs) and strong insulator signals, e.g. topological association domains (TADs) and the DNA binding sequence patterns of the CCCTC protein (CTCF binding motifs) (Bouvy-Liivrand et al. 2017; Suzuki et al. 2017). In mouse embryonic stem cells (mESCs), active enhancers that were bound by transcription factors Oct4, Sox2 and Nanog (OSN) (Whyte et al. 2013), and active promoters that were marked by H3K4me3 (Marson et al. 2008), were identified through ChIP-Seq. Interactions between promoters and enhancers in mESCs that were initialised by cohesin were identified by chromatin interaction analysis with paired-end tag sequencing (CHIA-PET) (Dowen et al. 2014). These data from mESCs showed that miRNA genes that were transcribed from genomic regions close to one another on a chromosome in mESCs were spatially close to a small subset of super-enhancers specific to mESCs (Suzuki et al. 2017). Super-enhancers are enhancers close to one another on the same DNA strand (typically ≤ 30Kb), which have been shown to control cellular identity (Khan and Zhang 2016). When CRISPR/Cas9 deletion was performed on each of the constituents of the miRNAs-associated super-enhancers, substantial but different levels of decrease in the *de novo* production of mature miRNAs were observed (Suzuki et al. 2017). Additionally, strong insulator signals from TAD boundaries and CTCF binding sites were observed between the transcription start sites (TSSs) of miRNAs close to one another on the chromosome (Bouvy-Liivrand et al. 2017). Those insulator elements separated the TSSs of miRNAs into two groups: miRNAs that were on the same side of the insulator elements as the super-enhancer expressed a large number of transcripts, whereas miRNAs that were on a different side of the insulator elements from the super­enhancer expressed fewer transcripts (Bouvy-Liivrand et al. 2017). These results indicated that the observation of identical miRNA sequences from different locations being expressed at different magnitudes in different bovine tissues could be because those miRNAs were controlled by different enhancers (or different constituents of the same super-enhancer), and/or because those miRNAs were separated by insulator elements, leading to only a subset of miRNAs accessible to enhancer regulatory control.

To predict the seed match sites of all expressed miRNAs on bovine mRNA transcripts, we firstly grouped expressed mature miRNAs into miRNA families based on their extended seed sequence (2^nd^ to 8^th^ nucleotides at the 5’-end of mature miRNA sequence). We found that 52% of novel mature miRNAs were grouped into the same miRNA families with mammalian miRNAs in miRBase (Kozomara and Griffiths-Jones 2014), indicating that these novel miRNAs had a similar function as known mammalian miRNAs in the same family. The remaining 48% of novel mature miRNAs did not group with any known miRNAs in miRBase. Along with those newly-discovered-known miRNAs that could not be grouped with any other miRNAs, the function of these miRNAs from recent discoveries could be inferred from the function of cognate targets that we predicted (Supplemental Table S4).

The 600 miRNA families that were formed from our expressed miRNAs were predicted to target 780,481 seed match sites on bovine mRNAs homologous to human. Because a putative seed match site could be shared between 1 and 3 miRNA families, there were 879,240 putative interactions between miRNA families and their mRNA targets (Supplemental Table S4). We demonstrated an enrichment of these putative interactions in interactions identified previously (Table 6). The fold change of enrichment, which compared the number of confirmed actual interactions with the average number of confirmed random interactions, were lower than 1.7 across all confirmation sets (i.e. miRTarBase, CLIP-Seq, miRWalk and TargetScan). This was because a miRNA family could interact with 4 to 6,767 putative target genes, which caused an incompletely shuffled dataset that had more permutated interactions identical to actual interactions and led to a lower level of fold change of enrichment.

*In-silico* identification of seed match sites on bostau6 was challenging because firstly, the prediction relied on the extended seed sequence within the mature miRNA sequence. miRDeep2 named miRNA sequences by comparing the precursor miRNA sequence from RNA-Seq data with that from miRBase (Friedlander et al. 2012). Mature miRNA sequences that were identified through this procedure could have the same name as that in miRBase even though the mature miRNA sequence, or even the seed sequence, differed by a few nucleotides from that in miRBase. This gave rise to the different putative seed match sites between miRNA and mRNA that were identified by TargetScan 7 (Agarwal et al. 2015) and by our study. Secondly, whole genome sequence (WGS) alignment from 100 vertebrates were required to filter false positive putative miRNA seed match sites (Lewis et al. 2005; Agarwal et al. 2015). Currently only mRNAs orthologous to humans were included in the WGS alignment, which was constructed from old versions of reference genomes including hgl9 and bostau7 (UCSC IDs) (Agarwal et al. 2015). As a result, miRNA seed match sites on bovine-specific mRNAs or on newly annotated mRNAs could not be identified using this method. Many miRNA seed match sites in bostau7 were not found in bostau6 because the human genes could not be identified through orthologs in bovine or because sequence compositions of the same gene were different between bostau7 and bostau6.

Confirmation of interacting miRNA sequences and mRNA sequences was also challenging because most public repositories only provided miRNA name and cognate target gene names. This made it difficult to determine (1) the strand of the DNA duplex where the mature miRNA was derived from, (2) the target gene names, because some were missing, and neither gene IDs or transcript IDs were provided, and (3) the target mRNA sequences, particularly when multiple target sequences were on the same target gene. To overcome these challenges, future studies should use CLIP-Seq to directly identify the interacting miRNA and target RNA target sequences. This will expand the repository of miRNA targets to all RNA transcripts. This will also validate the novel miRNAs, putative seed match sites and putative interactions that were identified in this study.

The choice of which strand of the duplex becomes the mature miRNA depends on the orientation of the duplex as well as the sequence context of the strand, but the mechanism is not fully known yet (Bartel 2018). Although most miRNAs only have one dominant strand consistent across samples, some miRNAs had both strands as the dominant products, while other miRNAs had different preferences for the strand between tissues, species or developmental stages (Griffiths-Jones et al. 2011; Choo et al. 2014). This observation has been called ‘arm switching’ or ‘arm selection’ (Griffiths-Jones et al. 2011). Examination of arm switching requires a large number of samples, and therefore in previous studies arm switching was more often examined across species (Griffiths-Jones et al. 2011; Kozomara and Griffiths-Jones 2011). With high throughput sequencing, arm switching has increasingly been shown between tissues or between cells (Kuchenbauer et al. 2011; Li et al. 2012; Gong et al. 2014; Kuo et al. 2015; Tsai et al. 2016; Lin et al. 2018), and was suggested to provide a fundamental mechanism to evolve the function of a miRNA locus and target gene network (Griffiths-Jones et al. 2011). In this study, we showed that cattle also had different arm usages in different tissues (Supplemental Table S2), and mature miRNAs that were produced from different strands of the same precursor miRNA led to different confirmed gene targets (Supplemental Table S4), similar to the different arm usages observed in humans, sheep, insects and rice (Marco et al. 2010; Griffiths-Jones et al. 2011; Bortoluzzi et al. 2012; Li et al. 2012; Hu et al. 2014; Kuo et al. 2015; Lagana et al. 2015; Tsai et al. 2016; Lin et al. 2018). Variants that control arm preference were suggested to be within the primary miRNA but outside the duplex (Griffiths-Jones et al. 2011). To identify the complete primary miRNA sequences, one could use nascent RNA assays such as CAGE (Kodzius et al. 2006) or PRO-Seq (Kwak et al. 2013) to identify the transcription start sites of RNA transcripts, use polyadenylated RNA termini assays such as 3P-seq (Jan et al. 2010) or TAIL-Seq (Chang et al. 2014) to identify the transcription termination sites, and use long-read RNA sequencing methods (Tilgner et al. 2015) to define the exact transcript in genomic regions with large repeats or large gene families.

Using 65 million WGS variants from the 1000 Bull Genomes Project (Daetwyler et al. 2014) (*Bos Taurus* Run 6), we showed that compared with the entire genome and mRNA transcripts, miRNA genes were depleted for common variants (Table 7), but were enriched for sequence variants with extreme allele frequencies (Table 8 Figure 4; Figure 5), supporting the hypothesis that miRNA genes are under strong natural selection. We also showed that miRNA target sites were more depleted for filtered variants than the entire genome but were less depleted than miRNA genes (Table 7). Additionally, miRNA target sites were more enriched for variants with extreme frequencies than the entire genome but were less enriched than miRNA genes (Table 8 Figure 4). These results indicate that miRNA target sites were under weaker natural selection than miRNA genes.

The stronger selection of miRNA genes and weaker selection of miRNA target sites could be a mechanism to retain the specific role that miRNA plays in the cellular processes yet also to contribute to the variation of phenotypes. To retain the specific functional role, miRNA genes are highly conserved in evolution, particularly at the 5’end of the mature miRNAs (Lee et al. 1993; Reinhart et al. 2000; Bartel 2018). Mutations in miRNA genes were expected to rewire the miRNA targeting network, affect many cellular processes and cause deleterious consequences to the animal. To compensate these strong effects, miRNAs expanded their members forming a miRNA family. If one miRNA gene was mutated, other members in the same family could be expressed to target the same transcript through the same seed target site (Agarwal et al. 2015; Bartel 2018). Alternatively, miRNAs from other families could target the same transcript through a different seed target site because there were often more than one target sites on a transcript (Supplemental Table S4). A mechanism for miRNAs to contribute to variation of phenotypes could be to mutate miRNA seed match sites and target sequences more frequently such as through insertions and deletions (INDELs) (Figure 5). Mutations in miRNA target sequences abolish or decrease the repression effects of cognate mature miRNAs. It has been suggested that each miRNA target site contributes a modest repression effect in most cases (with multiple target sites in the same 3’UTR adding up to much more substantial repression) (Baek et al. 2008; Selbach et al. 2008). By mutating miRNA target sequences, miRNAs could provide a finer tuning of the protein profile. Previous studies on the functions of miRNAs proposed that occasionally the collective function of a group of miRNAs could be enough to either trigger or sharpen a developmental transition, but more generally, miRNAs were expected to produce a much more complex topology of gene expression in the nucleus, with more optimal levels of protein synthesis in the cytoplasm of each cell of each tissue (Bartel 2018).

miRNAs have been found to have functional applications in cattle. *Bta-miR-103-2, bta-miR-150* and *bta-miR-181b-2* were shown to be up-regulated during heat events in Frieswal (*Bos Taurus* x *Bos Indicus*) crossbred dairy cattle (Sengar et al. 2018). Tissue collection for this cow was conducted in the early spring of Victoria, Australia when the weather was mild (Chamberlain et al. 2015), and these miRNAs were lowly expressed in this *Bos Taurus* dairy cow’s skin tissues (mature miRNA read count: 4-74). Whereas *bta-miR-142*, which has been shown to be down-regulated during heat event (Sengar et al. 2018), was moderately expressed in skin tissues (mature miRNA read count: 248-317). This provides the possibility of using these miRNAs as biomarkers for heat tolerant cattle.

The *DGAT1* gene is known to have a large effect on milk yield and composition in dairy cattle (Grisart et al. 2004).

In another study, a genomic region around *TFCP2* was associated with fertility in both Australian and Irish dairy cattle (Moore et al. 2016). We found 4 single-base-nucleotide substitutions, 0 insertion and 0 deletion from WGS variants from 1000 Bull Genomes Project (Daetwyler et al. 2014) (*Bos Taurus* Run 6) within the seed match sites within the *TFCP2* mRNA transcript (Table 9). The seed match site that had two WGS variants, *chr5:28806521-28806526(+)*, was predicted to be targeted by the *bta-miR-11995-5p* miRNA family. *TFCP2* is known to be targeted by *bta-miR-660-5p* (Chou et al. 2018), but in our data, there was no variant within the only putative seed match site, *chr5:73427696-73427701(+)*, for the *bta-miR-660-5p* family within the *TFCP2* mRNA transcript region. Both *TFCP2* and *bta-miR-660-5p* were expressed in all 17 tissues from this cow, whereas *bta-miR-11995-5p* was only detected in brain caudal lobe and brain cerebellum. Our prediction showed that the *bta-miR-11995-5p* miRNA family would have a higher repression effect than the *bta-miR-660-5p* miRNA family (Table 9), suggesting that in brain caudal lobe and brain cerebellum tissues, *bta-miR-11995-5p* was likely to bind to the *TFCP2* mRNA transcript to repress *TFCP2* translation.

**Table 9.**
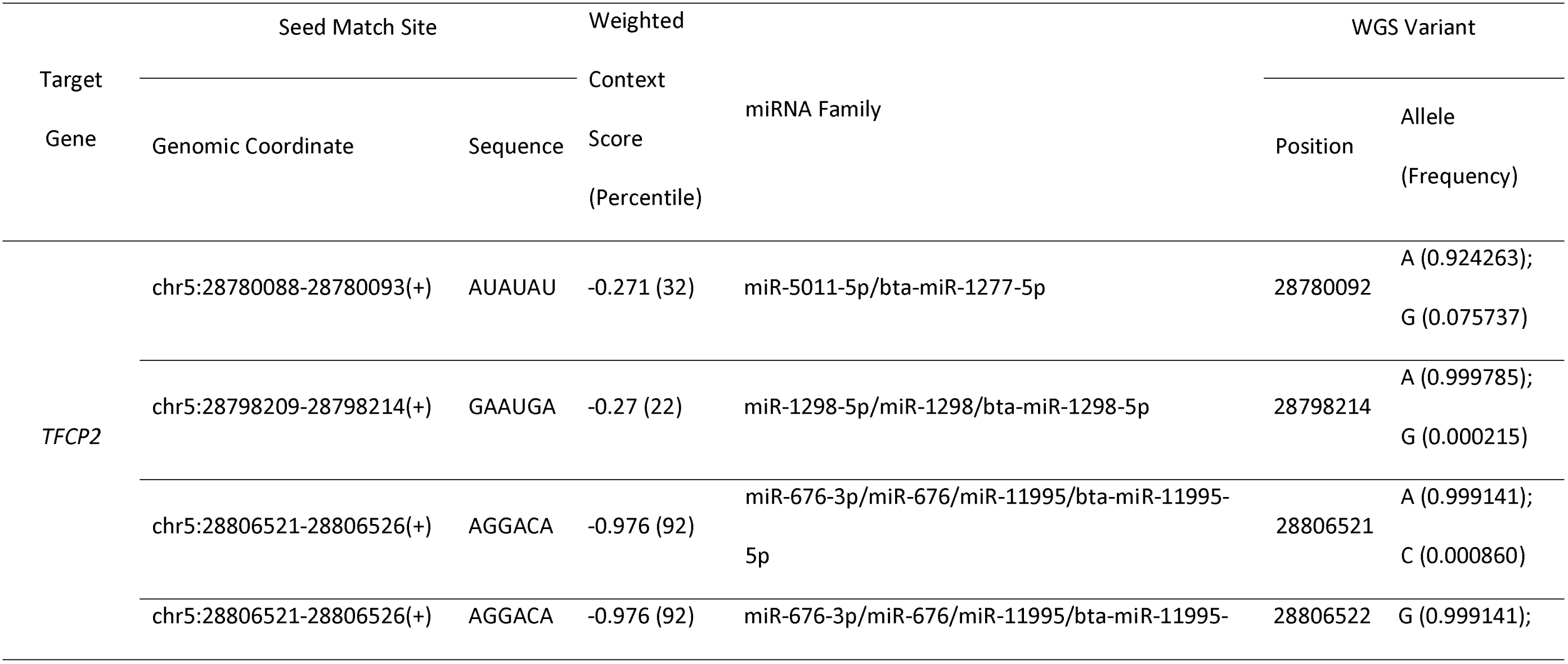
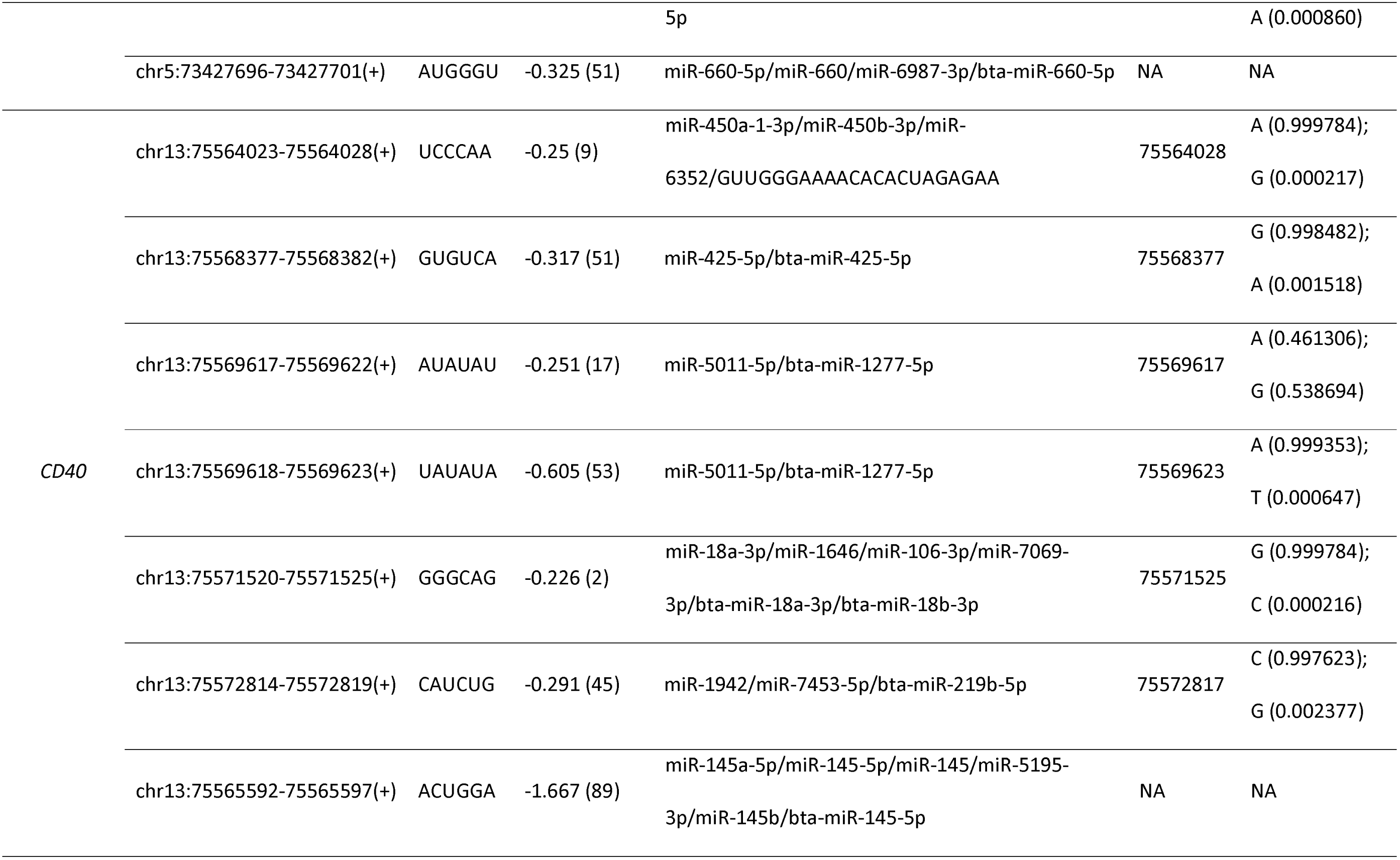
1000 Bull Raw WGS Variants within Seed Match Sites within *TFCP2* and *CD40 mRNA* Transcripts. Weighted context score and weight context score percentiles are statistics from TargetScan (Agarwal et al. 2015) prediction. The lower the weighted context score, the higher the predicted repression effect. The higher the weight context score percentile, the higher the confidence of the prediction. miRNAs are grouped into families and represented by multiple miRNA names connected by the forward slash symbol. miRNA names with a prefix of “miR-” are annotated miRNAs from miRBase. miRNA names with a prefix of “bta-miR-” are expressed known miRNAs in bovine. The rest are expressed novel miRNA sequence detected.

Similarly, gene *CD40* encodes the protein CD40 which is a receptor on antigen-presenting cells of the immune system and is essential for mediating a broad variety of immune and inflammatory responses including mastitis in dairy cattle (Lutzow et al. 2008). *CD40* is known to be regulated by *bta-miR-145-5p* (Chou et al. 2018). Our prediction showed that apart from *bta-miR-145-5p, CD40* could also be regulated by several other known or novel miRNAs, including the *miR-378g/miR-6637-3p/miR-7482-5p/GCUGGGCUGCGUCGGCGCUCGGA* family which had a more effective repression site than *bta-miR-145-5p* (Supplemental Table S4), indicating that these miRNAs are likely to target *CD40* if they are expressed. We found 6 single-base-nucleotide substitutions, 0 insertion and 0 deletion from WGS variants from 1000 Bull Genomes Project (Daetwyler et al. 2014) (*Bos Taurus* Run 6) within the seed match sites within the *CD40* mRNA transcript region (Table 9). Although none of those seed match sites that had WGS variants had a higher repression effect than *miR-bta-145-5p*, an A-to-G substitution mutation at position *chr13:75569617* within the seed match site *chr13:75569617-75569622(+)* was interesting, because in most cases the allele that formed the seed match sequence had a frequency of over 99% (Table 9), but at *chr13:75569617*, the *’A’* allele that formed the seed match sequence had a frequency of 46.13% (Table 9), indicating that variant *chr13:75569617* is an old mutation, and animals from the 1000 Bull Genomes Project have adapted to and favoured the alternative allele ‘*G*’.

Given both miRNAs and mRNAs are transcribed by RNA polymerase II in the nucleus, transcription of miRNAs and that of their target mRNAs was speculated to be correlated. Our results didn’t appear to support this speculation, as we detected no general association between miRNAs and their putative mRNA targets that were differentially expressed from the same tissue from the same cow (Table 5). We also did not observe significant association between the expression of mature miRNAs and the allele-specific expression of putative miRNA target sequences from the same tissue. Neither did we find the heterozygous sites within putative miRNA target sequences more significantly associated with allelic imbalance of exons within the same target genes than the heterozygous sites within the gene regions other than exons and putative miRNA target sequences. Additionally, of a total of 902,172 putative pairs of miRNA and target mRNA, we found that 26 putative pairs had the miRNA gene overlapped the target mRNA gene on the same DNA strand. Another 26 putative pairs had the miRNA overlapped the target mRNA gene region but on the antisense strand. There were 32,466 putative pairs had the miRNA transcribed from the same chromosome as the target mRNA but outside the target mRNA region. And the remaining 869,609 pairs had the miRNA transcribed from a different chromosome from the target mRNA. Overall our observations suggested that miRNAs and their putative target mRNAs were more likely to scatter at different transcriptional units within the same nucleus (Iborra et al. 1996; Osborne et al. 2004) and therefore showed different transcription rates. Since miRNAs are known to repress the translation of target mRNAs, instead of comparing transcriptional levels between miRNAs and target mRNAs, it will be interesting for future studies to investigate whether the transcription levels of miRNAs are associated with the translation levels of target proteins, and whether quantitative trait loci (QTLs) for the translation levels of individual protein are enriched within miRNAs or miRNA target sequences.

In conclusion, our study has profiled expressed miRNAs and predicted their mRNA targets in 17 bovine tissues from a single lactating dairy cow. We have demonstrated that although miRNAs and seed match sites are depleted for common variants compared with entire genome, they are enriched for rarer variants, providing evidence for miRNA genes and target sequences under natural selection. Additionally, through switching the dominant miRNA sequence, a conserved precursor miRNA can regulate different target mRNAs in different bovine tissues, potentially contributing to the specialised function of each tissue. We also bring a closer connection between miRNAs and enhancer RNAs by demonstrating that miRNA seed match sites are significantly enriched at active enhancer regions. Contributing to the goal of the Functional Annotation of Animal Genomes (FAANG) consortium, our results help to complete the catalogue of tissue-specific interaction network between miRNAs and mRNA targets, which in the future may assist in better assessing the combined effects of genes and their regulators on dairy traits.

## Materials and Methods

All methods are summarized in a flow diagram provided in Supplemental Figure S1.

### Samples, RNA sequencing and alignment

Seventeen tissues from a dairy cow were collected in the early spring (8 September 2010) at Ellinbank, Victoria, Australia as part of another study (Chamberlain et al. 2015). The cow was 25 months old and 65 days into her first lactation, non-pregnant, and was euthanised following an incurable injury.

Small (10-50 nucleotides) RNA transcripts in the cell nucleus and cytosol were isolated from 50mg of ground frozen tissue using the mirVana miRNA isolation kit (Ambion) in duplicate as per the manufacturer’s instructions.

Libraries were prepared for sequencing using the NEBNext Multiplex Small RNA Library Prep kit (New England Bioloabs Inc) according to manufacturer’s instructions with the following modification. Following PCR 6ul of each library were pooled and purified with the Qiaquick PCR purification kit (Qiagen) as per the manufacturer’s instructions. A second elution was performed for a total eluate of 60ul. 45ul of purified library pool was run with 6X loading buffer on a 3% TBE gel in 2 lanes alongside 5ul of Quick Load pBR322 DNA MspI Digest. The gel was run at 70 volts for 2 hours. Short RNA libraries were cut from the gel at 130-170bp with an expected insert size of 10-50nt. DNA was extracted from gel slices using the Qiagen QIAEX II gel extraction kit (Qiagen) as per the manufacturer’s instructions and eluted in 40ul Tris.

Libraries were sequenced on HiSeq3000 (Illumina) in a paired-end 50 cycle run and FASTQ files were generated with bcl2fastq2 conversion software. Raw sequence quality was assessed using FastQC (Andrews 2010) (version 0.10.1).

Illumina adapter sequences and poor-quality bases were trimmed from raw RNA sequence reads. Two trimming pipelines were used and compared. The first was cutadapt (Martin 2011) (version 1.9) and sickle (Joshi and Fass 2011) (version 1.33) following the micro RNA-Seq Data Processing Pipeline from Encyclopedia of DNA Elements (ENCODE) consortium. Raw RNA sequence reads were trimmed from the 3’-end when ≥5 consecutive base pairs matched with the adapter sequences. Trimmed sequence reads were discarded if final read length was <18nt or had low quality score (read error rate >0.1). The second, trimmomatic (version 3.6) (Bolger et al. 2014), trimmed raw RNA sequence reads from the 3’-end when ≤2 nucleotides matched with the adapter sequence during the standard ‘seed and extend’ approach (Li and Homer 2010) and ≥5 consecutive nucleotides matched with the whole adapter sequence. To trim partial read-through adapter sequences, initial-trimmed RNA sequence reads were trimmed from the 3’-end when >10 consecutive nucleotides matched with part of the adapter sequence. Nucleotides at both ends were further trimmed off if read quality was <15. The remaining reads were trimmed from the 3’-end if the averaged read quality among 4 consecutive nucleotides were <15. Trimmed reads were retained if final read length was ≥18nt. Trimmed sequence quality for all trimmers was also assessed using FastQC (Andrews 2010) (version 0.10.1).

Trimmed paired sequence reads were aligned to bovine reference genome UMD3.1.1. (bostau8) using BWA (Li and Durbin 2009) backtrack algorithm (aln) (version 0.7.17), bowtie (Langmead 2010) (version 1.2.2), bowtie2 (Langmead and Salzberg 2012) (version 2.3.4.1), STAR (Dobin et al. 2013) (version 2.5) and HISAT2 (Kim et al. 2015) (version 2.1.0) respectively. Default settings were applied to all aligners, except that the maximum insert size was 50nt in BWA, the overhang was 49 (maximum sequence length minus 1) (Dobin et al. 2013) in the first step of the two-step STAR alignment, and both mapped and unmapped reads were required to return in the output BAM file for all aligners. Alignment statistics were generated using SAMtools (Li et al. 2009) (version 1.6). Only mapped and paired reads with mapping quality over 20 were retained for analysis.

### Identification of expressed known and novel miRNAs

Canonical and non-canonical micro RNAs (miRNAs) that were expressed in 17 bovine tissues were identified using miRDeep2 (Friedländer et al. 2012) (version 2.0.0.8). Aligned paired RNA sequence reads with mapping quality ≥20 were provided as input to the second module of miRDeep2 (Quantifier; default settings) and processed through to the last module (miRDeep2; default settings). The first module (Mapper) was excluded because it required raw single-end RNA sequence data as input, and the built-in trimmer did not take partial adapter read-through into account. Technical replicates were provided as input to the miRDeep2 pipeline. Also provided were the complete collection of precursor and mature miRNA sequences from bovine and bovine-related species from miRBase (Kozomara and Griffiths-Jones 2014) (version 22), where goat and sheep were chosen as the bovine-related species. A surge of miRNAs from deep sequencing had been added to the complete collection of miRBase (version 22), but the high confidence set from miRBase (version 22) had not been updated at the time this study was conducted and therefore was not used. Consistent with the case study in the miRDeep2 paper (Friedländer et al. 2012), candidate miRNAs that resembled other small RNAs such as tRNA and rRNA were removed. The miRDeep2 algorithm assigns each novel precursor miRNA a log-odds score, which indicates the probability that the sequence is a true miRNA precursor instead of a background hairpin (Friedländer et al. 2008; Friedländer et al. 2012). For each analysis, the lowest miRDeep2 score cut-off that yielded a signal-to-noise ratio of 10:1 or higher was used to select the candidate miRNAs for further analysis. The actual threshold for each library is listed in “filters for each library” tab of Supplemental Table S2.

miRDeep2 returned the genomic coordinates of precursor miRNAs and the DNA sequence aligning to cognate mature and star miRNAs (Friedländer et al. 2012). The genomic coordinates of mature and star miRNAs were identified by searching respective sequences within the cognate precursor. ‘-5p’ and ‘-3p’ ends were identified by comparing the relative genomic positions of mature and star miRNAs that were derived from the same precursor. Due to variation among sequence libraries, a stack of sequence reads aligning to the same known precursor, mature or star miRNA varied in length (e.g. Figure 2) (Friedländer et al. 2012). In those cases, a consensus sequence was derived from the stack of reads, and the genomic coordinates were updated accordingly. The consensus sequence was selected as follows:

1. If the stack of reads were classified as known and fell into a miRNA region annotated in UMD3.1.1 (GCF_000003055.6), the consensus sequence was the sequence in UMD3.1.1 annotation.
2. If the stack of reads fell outside the annotated miRNA regions in UMD3.1.1, the consensus sequence was the sequence with the highest miRDeep2 score (Friedländer et al. 2012). miRDeep2 scores a miRNA by considering the hairpin folding, the mature, star and loop sequence from the hairpin, and the proportion of nucleotides in the mature miRNA passing the RNA folding threshold from the randfold software (Bonnet et al. 2004; Lorenz et al. 2011) (Github: ebOO/randfold).
3. If multiple sequences reached the same highest miRDeep2 score, the consensus sequence was the sequence with the highest frequency across all libraries.
4. If multiple sequences reached the same highest frequency, the consensus sequence was the longest sequence.
5. If multiple sequences reached the same longest length, the consensus sequence was the collapsed sequence from the stack (this only applies to mature and star sequences).

The consensus miRNA genomic coordinates and miRNA sequences were considered as “clean” miRNAs and were retained for analysis. Note, the observation that a stack of reads aligning to the same known precursor, mature or star miRNA varied in length in this lactating dairy cow, is similar to the miRNA isoforms that were detected in healthy humans, where depending on gender, population and race, precursor miRNAs gave rise to many isoforms that typically differed in either 5’ or 3’ termini or both, and the most abundant isoform was frequently annotated as the canonical sequence in miRBase (Telonis et al. 2015; Magee et al. 2018). However, miRNA isoform was out of the scope of this study, and our method to obtain consensus sequence didn’t consider miRNA isoforms.

### Identification of differentially expressed mature miRNAs

Mature miRNA read counts from miRDeep2 were used to produce a tissue-by-miRNA read count matrix (17 tissues x 610 miRNAs). The final read count of a miRNA was the averaged read count between technical replicates. miRNAs with <10 reads across all tissues were removed from the analysis. DESeq2 (version 1.18.1) (Love et al. 2014) with default shrinkage estimator was used to identify mature miRNAs that were more or less often expressed in a tissue than the average of all other tissues. A miRNA was considered differentially expressed if the miRNA was found expressed by miRDeep2, and after adjustment for multiple testing (Benjamini-Hochberg correction) the P-value was <0.01, and the expression of miRNA in the tissue was >2 fold different to the averaged expression across all other tissues.

### Prediction of miRNA seed match sites

TargetScan suite (Lewis et al. 2005; Friedman et al. 2009; Agarwal et al. 2015) (version 7.2) was used to predict the seed match sites of expressed miRNAs. All input files and Perl scripts from TargetScan 7 website (downloaded on 19 June 2018) were used in our analysis except the miRNA seed file, miRNA sequence file and the UTR profile file were prepared as per instruction from TargetScan 7 (Agarwal et al. 2015).

The miRNA seed file was table of grouped miRNA names (i.e. miRNA family), seed sequence, and grouped animal IDs from miRBase that contributed constituents of the miRNA family. To construct the miRNA file, all cleaned mature miRNA sequences from 17 bovine tissues, and the mature miRNA sequences of ten vertebrates from miRBase (Kozomara and Griffiths-Jones 2014) (version 22), were grouped into miRNA families based on the identity of extended seed sequence (2^nd^-8^th^ nucleotides from 5’-end of mature miRNA). The ten vertebrates from miRBase were the same as those in TargetScan 7 predictions, which were mouse, rat, opossum, western clawed frog, chicken, rhesus macaque, chimpanzee, human, dog and cattle.

The miRNA sequence file was a table of grouped miRNA names (i.e. miRNA family), animal ID from miRBase that contributed constituents of the miRNA family, miRBase ID of a constituent of the miRNA family and the mature miRNA sequence. miRNA family ID, miRBase species ID, miRBase ID and mature miRNA sequence were sorted according to the miRNA file.

The UTR profile file was a table of transcript ID or gene ID or gene name from human reference genome hg19 (UCSC ID), animal ID from miRBase that contributed the 3’UTR sequence for whole genome sequence (WGS) alignment, and the aligned 3’UTR sequence. As per instruction from TargetScan 7 (Agarwal et al. 2015), the UTR profile was constructed from WGS alignment of 84 vertebrate species.

TargetScan 7 did not directly output the genomic coordinates of putative miRNA seed match sites on to UMD3.1 (NBCI) or bostau6 (UCSC), but instead provided the positions of the seed match sites on the UTR profile. To identify the genomic coordinates of putative miRNA seed match sites on bostau6 chromosome 1-29, X and mitochondria, the Ensembl transcript IDs on bostau6 that were orthologs to hgl9 (UCSC ID) were identified using the getLDS function from R Bioconductor biomaRt package (version 2.34.2). Then the genomic coordinate of each seed match site on bostau6 was identified through local pairwise alignment between the miRNA seed sequence and the bostau6 gene sequence, using the pairwiseAlignment function (parameter: type=’local’) from R Bioconductor Biostrings package (version 2.46.0).

### Confirmation of putative interactions between miRNAs and targets

High-throughput sequencing of RNAs isolated by crosslinking immunoprecipitation (HITS-CLIP, also known as CLIP-Seq) was used to identify miRNA and mRNA sequences that were bound by the Argonaute (AGO) protein in bovine kidney cells (Scheel et al. 2017). The data was a table (S3) listing target gene name, genomic coordinate of target sequence (i.e. chromosome, start, end and strand), target sequence and cognate miRNA name. The miRNA target sequences were aligned to bovine reference genome Btau4.6.1 (NCBI) or bostau7 (UCSC). We converted their genomic coordinates to UMD3.1 (NCBI) or bostau6 (UCSC) using the UCSC liftOver tool (Kuhn et al. 2013).

To confirm putative interactions between bovine miRNA families and their cognate seed match sites using the Scheel *et al*. set, we identified putative miRNA seed match sites for miRNAs that were expressed in kidney tissues and also overlapped with miRNA target sequences in the Scheel *et al*. set. Then for each putative seed match site, we examined whether the miRNA name in the Scheel *et al*. set could be found in the miRNA family in the prediction set. If the miRNA name in the Scheel *et al*. set was found, the putative interaction between the miRNA seed match sites and cognate miRNA was considered “confirmed”.

miRTarBase (Chou et al. 2018) (version 7.0) is a public database that used natural language processing techniques to collect experimentally identified “miRNA and cognate gene target” interactions in animals from published literature. The animals were *Bos Taurus, Caenorhabditis elegans, Canis familiaris, Drosophila melanogaster, Danio rerio, Gallus Gallus, Homo sapiens, Mus musculus, Rattus norvegicus, Sus scrofa, Xenopus tropicalis* and *Ovis aries*. The experimental identification techniques included but not limited to qRT-PCR, Luciferase reporter assay, western blot, microarray and CLIP-Seq. The data consisted of a table listing miRTarBase ID, miRNA name, species, target gene name, experimental technique and PubMed ID. Gene names from species other than bovine were converted to orthologous bovine gene names using the getLDS function from R Bioconductor biomaRt package (version 2.34.2). Any records with target gene name as NA were removed from analysis.

TargetScan (Agarwal et al. 2015) (version 7.0) published putative miRNA seed match sites from human reference genome GRCh37 (NCBI) or hg19 (UCSC) for all miRNAs in miRBase (version 21). The data consisted of a table listing the genomic coordinate of seed match site, gene target and cognate miRNA family. Human target gene names were converted to orthologous bovine gene names using the getLDS function from R Bioconductor biomaRt package (version 2.34.2). Genomic coordinates of human miRNA seed match sites were converted to bovine reference genome bostau6 using the UCSC liftOver tool (Kuhn et al. 2013). Any records with target gene name as NA were removed from analysis.

miRWalk (Dweep and Gretz 2015) (version 3.0) is a public database for computationally-predicted interactions between miRNAs and cognate gene targets from human, mouse, rat, cow and dog using a random-forest-based approach. We downloaded the cow putative interactions set. The data consisted of a table listing miRNA name, target transcript Ensembl ID and target gene name.

To confirm putative interactions between miRNA families and cognate targets using miRTarBase, TargetScan and miRWalk, target gene names in miRTarBase and TargetScan and target transcript IDs in miRWalk (Dweep and Gretz 2015) were matched exactly with that in the our predicted miRNA targets. Then for each record, the cognate miRNA name in the miRTarBase, TargetScan or miRWalk was searched within the miRNA family in our prediction result. If the miRNA name was found, the putative interaction between the miRNA family and cognate targets was considered “confirmed”.

Messenger RNAs, which were either up- or down-regulated in one tissue compared with the mean expression in the other 17 tissues from the same lactating cow as used in this study were identified (Chamberlain et al. 2015). The tissue that was not included in our study was white blood cells. The data consisted of a table listing the Ensembl gene ID, gene name, gene genomic coordinates, the tissue where the gene was differentially expressed, fold change (all >2-fold change) and P-value after corrected for multiple testing (all <0.01). Also provided were summaries of the GO terms and KEGG pathways of those differentially expressed genes.

The differentially expressed genes (Chamberlain et al. 2015) were combined with the differentially expressed miRNAs that were identified in this study to create a differential expression set. To identify miRNAs and cognate putative gene targets that were differentially expressed in the same tissue of the same cow, differentially expressed gene IDs were matched exactly with putative target gene IDs. Then the tissues where the genes were differentially expressed in were matched exactly with the tissues where the miRNAs were differentially expressed. For each putative interaction between the miRNA and the mRNA within tissue, if the miRNA name in the differential expression set was found in the miRNA family in the prediction set, then the miRNA and cognate gene target were considered as differentially expressed in the same tissue of the same cow.

### Enrichment analysis

A permutation test was performed to examine whether putative interactions between the miRNA families and cognate targets (named as the ‘prediction set’) overlapped with interactions from previous experimental and computational identifications (named as the ‘confirmation set’) more often than by chance. Data from CLIP-Seq (Scheel et al. 2017), miRTarBase (Chou et al. 2018), TargetScan (Agarwal et al. 2015) and miRWalk (Dweep and Gretz 2015). The actual number of overlapping interactions between the prediction set and the confirmation set, which was denoted as n, was the same as was described in the previous section (Confirmation of putative interactions between miRNAs and targets). To create a random interaction set, the list of miRNA families in the prediction set was shuffled randomly and then combined with the list of predicted targets. The number of overlapping interactions between the random set and the confirmation set, which was denoted as m, was then calculated. This shuffling and counting procedure were repeated 10,000 times. The ranking position of *n* within the distribution of 10,000 *m* values, denoted as *R*, was determined, and a P-value to test the significance of the ranking was computed. If n was larger than all 10,000 m values, the P-value was set to < 0.0001 and otherwise it was 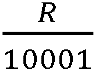. The fold change of enrichment was defined as the ratio of the actual number of validated pairs to the average number of validated pairs in 10,000 random sampling.

### Polymorphisms in miRNA genes and targets

A genomic feature is a predefined genomic range, which could be miRNAs that were expressed in one or more bovine tissues, miRNA targets that were identified through experimental or computational procedures, or the entire bovine genome (genome-wide). Raw (65,195,092) and filtered (Daetwyler et al. 2017) (44,678,426) WGS variants from 1000 Bull Genomes Project (Daetwyler et al. 2014) (*Bos Taurus* Run 6) from 2,333 individuals were utilised. Raw and filtered variants within a genomic feature were examined in the following three ways:

1. The polymorphic rate in a genomic feature, *y*_1_, was defined as follows:

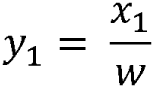

Where *x*_1_ was the total number of polymorphic sites (including single-base-nucleotide substitutions, insertions and deletions) within the genomic feature, and *w* was the total width of the same genomic feature. Here, the map file for the filtered WGS variants in TXT format was used to calculate *x*_1_.
2. The density of rarer variants with an allele frequency less than a in a genomic feature, y2, was defined as follows:

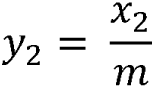

Where *x*_2_ was the number of variants (single-base-nucleotide substitutions only) with an allele frequency less than *α* within the genomic feature, and *m* was the total number of variants (single-base-nucleotide substitutions only) within the same genomic feature. Here, the variant call file for all raw sequence variants in VCF format was provided as input to the vcftools (version 0.1.15) to calculate the allele frequency for each variant, and custom R scripts were used to calculate *x*_2_ and *m*. If >1 alternative allele at a locus was detected, the frequency of each alternative allele was calculated.
3. INDELs were categorised by the number of positional shifts, *n ∊* (−∞, +∞), between the alternative allele and reference allele at the same locus, following a classification system similar to what was previously used (Mills et al. 2006; Bhattacharya et al. 2012). The density of INDELs with an n-th positional shift in a genomic feature, *y*_3_, was defined as follows:

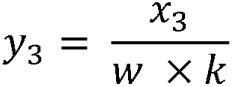

Where *x*_3_ was the number of variants (INDELs only) with an *n*-th positional shift comparing the alternative allele with the reference allele among all the *k* animals from the 1000 Bull Genomes Project (Daetwyler et al. 2014) (*Bos Taurus*, Run 6) and *w* was the total width of the same genomic feature. Here, the variant call file for all raw sequence variants in VCF format was provided as input to the vcftools (version 0.1.15) to count the number of alleles (INDELs only) at each locus, and custom R scripts were used to calculate *x*_3_, *w* and *k*. If >1 alternative allele at a locus was detected, each alternative allele was calculated.

### Examination of associations among miRNAs, miRNA target regions and allele-specific expression of target genes

RNA sequence (RNA-Seq) data was previously used to assess the degree of allelic imbalance across 18 tissues from the same cow (Chamberlain et al. 2015). The degree of allelic imbalance was estimated as a Chi-Square value at each heterozygous site that was identified from the 1000 Bull Genomes Project (Daetwyler et al. 2014) (*Bos Taurus* Run 5). Although only results within exons were published, allelic imbalance had been calculated for heterozygous sites within all genes that were annotated in UMD3.1 (Ensembl release 75). We obtained this ASE data (Chamberlain et al. 2015), and defined ASE scores as the square root of Chi-Square values from their ASE analysis.

Very few variants were located within putative miRNA seed match sites because putative miRNA seed match sites were short (6nt) and highly conserved (perfectly reverse-complementary to miRNA seed sequences). Because sequence contexts around miRNA seed match sites were shown to affect miRNAs binding to target mRNA sequences (Agarwal et al. 2015), we created putative miRNA target sequences by extending 49nt upstream and 1nt downstream of putative miRNA seed match sites. This would include the 2^nd^-7^th^ nucleotides within mature miRNA sequences perfectly matched with the putative miRNA target sequences, and the lengths of putative miRNA target sequences, which were all 56nt, equivalent to the maximum length of miRNA target sequences that was identified by a CLIP-Seq experiment (Scheel et al. 2017).

We asked the following two questions:

1. Across tissues, was the allelic imbalance at each heterozygous site within putative miRNA target sequence associated with the expression level of cognate mature miRNA?
2. Across genes, were genes with heterozygous sites within putative miRNA target sequences more likely to show allelic imbalance at exons than genes with homozygous sites within putative miRNA target sequences?

To answer question 1, we calculated ASE scores for 459,610 heterozygous sites in 15,091 genes on 30 chromosomes (1-29 and X) in this animal. We selected ASE scores from heterozygous sites within putative miRNA target sequences. This dataset was combined with the dataset of mature miRNA read counts from miRNAs that were expressed in the same tissue as where ASE scores were measured from.

Analysis of variance (ANOVA) was used to assess if the expression level of mature miRNAs had an effect on the allelic imbalance of putative miRNA target sequence. The model was:

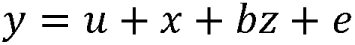

Where *y* was the ASE scores of heterozygous variants at putative miRNA target sequences, *u* was the mean, *x* was the effect of the variant names which was a factor variable, *b* was the coefficient for *z*; *z* was the read counts across tissues of a cognate miRNA, and *e* was the residue.

To answer question (2), all 30,831,575 polymorphic sites in the cow’s WGS variants from 1000 Bull Genomes (Daetwyler et al. 2014) (*Bos Taurus* Run 5) were used to identify polymorphic sites within all putative miRNA target sequences. A label of 0 and 1 was used to distinguish whether a polymorphic site was homozygous or heterozygous, respectively.

A linear regression model was used to assess whether zygosity within putative miRNA target sequences had an effect on allelic imbalance in exons of miRNA target genes. The model was:

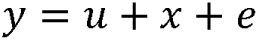

Where *y* was the ASE scores from heterozygous variants at exons of target genes, *u* was the mean, *x* was the effect of a factor variable that labelled whether a polymorphic site within a putative miRNA target sequence was homozygous or heterozygous in this cow, and *e* was the residue.

A permutation test was performed to examine whether the observed result in question (2) was due to linkage disequilibrium (LD) between variants within putative miRNA target sequences and that within exons. To construct the null miRNA target sequences, we selected genomic regions that were within putative miRNA target genes but were neither putative miRNA target sequences nor exons that were annotated in UMD3.1 (Ensembl release 91). Then we selected WGS variants from 1000 Bull Genomes (Daetwyler et al. 2014) (*Bos Taurus* Run 5) that were within null miRNA target sequences. We calculated the number of WGS variants that were within the original putative miRNA target sequences and defined this number as *N*. To construct the null dataset, we combined the dataset from randomly selecting *N* variants within null miRNA target sequences with the dataset of ASE scores from exonic heterozygous sites by the same genes. The null dataset was used to fit the same linear regression model as that in question (2). This procedure of constructing the null dataset and then fitting linear regression model was repeated 10,000 times. If the coefficient and standard error of coefficient from linear regression model that was constructed from the original dataset were respectively all bigger and all smaller than those from the 10,000 null datasets, we declared the observed result in question (2) was statistically significant.

## List of Abbreviations

ANOVA: An analysis of variance
ASE: Allele-specific expression
CAGE: Cap Analysis Gene Expression
CDS: coding sequence
CHIA-PET: chromatin interaction analysis with paired-end tag sequencing
CTCF: CCCTC binding factors
ChIP-Seq: chromatin immunoprecipitation followed by high-throughput sequencing
DE: Differential Expression
GWAS: Genome Wide Association Study
HITS-CLIP: high-throughput sequencing of RNAs isolated by crosslinking and immunoprecipitation, also known as CLIP-Seq
H3K27ac: acetylated lysine 27 on histone H3
H3K4me3: tri-methylation of lysine 4 on histone H3
INDELs: insertions or deletions
LD: linkage disequilibrium
PRO-Seq: precision run-on sequencing
QTLs: quantitative trait loci
ORF: open reading frame
RNA: ribonucleic acid
RNA-Seq: RNA Sequencing
SE: Super-enhancers
SINE: short interspersed elements
SNP: single nucleotide polymorphism
TAD: Topological Association Domains
WGS: whole genome sequence
mRNA: message ribonucleic acid
miRNA: micro ribonucleic acid
pre-miRNA: precursor micro ribonucleic acid
pri-miRNA: primary micro ribonucleic acid
5’UTR: five-prime untranslated region
3P-seq: poly(A)-position profiling by sequencing
3’UTR: three-prime untranslated region

## Declarations

### Ethics approval and consent to participate

No animal experiments were performed specifically for this work. The data was obtained from existing samples, references for these experiments are provided.

### Consent for publication

The authors agree for the publication of this manuscript to the journal RNA.

### Data availability

Micro RNA sequence data has been deposited under European Nucleotide Archive (ENA) accession number ERX2749848 - ERX2749881.

## Funding

The authors thank the DairyBio Initiative, which is jointly funded by Dairy Australia and Agriculture Victoria Research (Melbourne, Australia), for the generous funding for this project.

### Author’s contributions

MW performed all the data analysis and wrote the manuscript. MEG, TPH and BJH supervised the statistical analysis. CPP and AJC collected samples and sequenced RNA reads. CVJ provided data for the allele-specific expression analysis and differentially expression analysis for mRNAs from 18 tissues from this lactating dairy cow. MW, AJC, JEP, BGC, MEG and BJH conceived experimental design and coordination. All authors read, commented and approved the final manuscript.

## Acknowledgements

We acknowledge partners in the 1000 Bull Genomes Project for granting access to the whole genome sequence variants in the *Bos Taurus* genomes. We are immensely grateful to Daisy, the young dairy cow who was raised at the foothills of the rolling Strzelecki Ranges of West Gippsland, the National Centre for Dairy Research and Development at Ellinbank, Victoria, Australia.

## Supplementary Information

### Supplemental Table SI

#### Short RNA Sequence Reads, Filtering and Alignment

Trimming summary is given for each sequence library (tissue, technical replicate), a tag of whether Illumina adapter combination was detected in raw sequence, in trimmed sequences from cutadapt (Martin 2011) and sickle (Joshi and Fass 2011), and in trimmed sequences from trimmomatic (Bolger et al. 2014) (Illumina adapter contamination detected), total number of reads in raw and trimmed libraries (total number of reads), read length of raw and trimmed reads (read length), and proportion of guanine and cytosine nucleotides presented in raw and trimmed reads (GC ratio).

Alignment summary is given for each sequence library (tissue, technical replicate), total number of reads input to aligner (Total number of reads), number of reads from each end (Read 1, Read 2), number and ratio of mapped reads (Mapped reads), number of unmapped reads (Unmapped reads), number of paired reads mapped (Reads mapped and paired), reads with a mapping quality of 0 (reads mapped and MQ=0), inferred averaged insert size (insert size averaged), paired reads in inwards orientation (inward orientated reads) and all other alignment statistics from samtools (Li et al. 2009) stats option (version 1.6).

### Supplemental Table S2

#### Known and Novel miRNAs Expressed in 17 Bovine Tissues

A table for all expressed known and novel miRNAs that were detected by miRDeep2 (Friedländer et al. 2012). Significance threshold was the lowest miRDeep2 score that yielded a signal-to-noise ≥10:1 in each library, and miRNAs that passed significance threshold were retained for analysis. miRNA type specified whether the record was from a precursor, mature or star miRNA. Records with the same rowIndex indicated the mature and star miRNAs were derived from the precursor miRNA. Also presented are sequence library (tissue, technical replicate), genomic coordinates of the expressed miRNA (chromosome, start, end, strand, Refseq), miRNA duplex origin (miRNA name), miRNA sequence, information from miRBase (Kozomara and Griffiths-Jones 2014) (version 22) (miRDeep2 type, miRBase ID), confidence of the detection (miRDeep2 score, estimated probability of true positives, total read count, mature read count, star read count), whether arm switching was detected (armSwitch), and whether the known miRNA sequence has been updated to align with the annotation in UMD3.1.1.

### Supplemental Table S3

#### Differentially Expressed Mature miRNAs

Presented is results of differentially expression analysis using DESeq2 (Love et al. 2014). Each mature
miRNA with a sum read count across all tissues ≥10 was assessed whether the mature miRNA (chr, start, end, strand, miRNA name, miRNA sequence) was more often expressed in a lactating cow’s tissue than the mean expression of all other tissues. The result was presented with P-value after correction for multiple testing and the logarithm with a base of 2 of the fold change of expression.

### Supplemental Table S4

#### Predict miRNA Seed Match Sites

Presented is putative miRNA seed match sites that were predicted using TargetScan (Agarwal et al. 2015). Provided were the miRNA seed match sites (chr_target, seedMatch_start, seedMatch_end, strand_target) on bovine protein coding transcripts (Ensembl transcriptld_target, geneld_target, geneName_target), and the cognate miRNA_family.

### Supplemental Table S5

#### Examples of Different miRNA Arm Usages Leading to Different mRNA Targets

Presented is confirmed putative interactions between expressed miRNA and cognate gene targets by miRTarBase (Chou et al. 2018) and CLIP-Seq (Scheel et al. 2017).

### Supplemental Figure S1

#### Flow Diagram Depicting Steps Taken in This Study

Presented is a flow diagram of the process to identify expressed miRNAs from deep sequencing
samples from 17 tissues of a lactating dairy cow using miRDeep2, and to identify putative miRNA seed match sites of all expressed miRNAs using TargetScan. Differential expression analysis and enrichment analysis were also performed to confirm putative interactions between expressed miRNAs and cognate targets aligning with known interactions from public repositories before the examination of polymorphic sites within miRNA genes and targets. Colours were used to separate each analysis, as well as datasets, software and processing steps involved.

## Reference

Abelson JF, Kwan KY, O’Roak BJ, Baek DY, Stillman AA, Morgan TM, Mathews CA, Pauls DL, Rašin M-R, Gunel M et al. 2005. Sequence Variants in *SLITRKl* Are Associated with Tourette’s Syndrome. Science 310: 317–320.

Agarwal V, Bell GW, Nam J-W, Bartel DP. 2015. Predicting effective microRNA target sites in mammalian mRNAs. eLifeA: e05005.

Altuvia Y, Landgraf P, Lithwick G, Elefant N, Pfeffer S, Aravin A, Brownstein MJ, Tuschl T, Margalit H. 2005. Clustering and conservation patterns of human microRNAs. Nucleic Acids Research 33: 2697–2706.

Andersson L, Archibald AL, Bottema CD, Brauning R, Burgess SC, Burt DW, Casas E, Cheng HH, Clarke L, Couldrey C et al. 2015. Coordinated international action to accelerate genome-to-phenome with FAANG, the Functional Annotation of Animal Genomes project. Genome Biology 16: 57.

Andrews S. 2010. FastQC: a quality control tool for high throughput sequence data. *Available online at:* http://wwwbioinformaticsbabrahamacuk/proiects/fastgc.

Baek D, Villén J, Shin C, Camargo FD, Gygi SP, Bartel DP. 2008. The impact of microRNAs on protein output. Nature 455.

Bartel DP. 2018. Metazoan MicroRNAs. Cell 173: 20–51.

Bender W. 2008. MicroRNAs in the Drosophila bithorax complex. Genes & Development 22: 14–19.

Betel D, Koppal A, Agius P, Sander C, Leslie C. 2010. Comprehensive modeling of microRNA targets predicts functional non-conserved and non-canonical sites. Genome Biology 11: R90.

Bhattacharya A, Ziebarth JD, Cui Y. 2012. Systematic Analysis of microRNA Targeting Impacted by Small Insertions and Deletions in Human Genome. PLOS ONE 7: e46176.

Bolger AM, Lohse M, Usadel B. 2014. Trimmomatic: a flexible trimmer for Illumina sequence data. Bioinformatics 30: 2114–2120.

Bonnet E, Wuyts J, Rouzé P, Van de Peer Y. 2004. Evidence that microRNA precursors, unlike other non-coding RNAs, have lower folding free energies than random sequences. Bioinformatics 20: 2911–2917.

Bortoluzzi S, Bisognin A, Biasiolo M, Guglielmelli P, Biamonte F, Norfo R, Manfredini R, Vannucchi AM. 2012. Characterization and discovery of novel miRNAs and moRNAs in *JAK2*V617F-mutated SET2 cells. Blood 119: el20–el30.

Bouvy-Liivrand M, Hernández de Sande A, Pölönen P, Mehtonen J, Vuorenmaa T, Niskanen H, Sinkkonen L, Kaikkonen MU, Heinäniemi M. 2017. Analysis of primary microRNA loci from nascent transcriptomes reveals regulatory domains governed by chromatin architecture. Nucleic Acids Research 45: 9837–9849.

Budak H, Bulut R, Kantar M, Alptekin B. 2016. MicroRNA nomenclature and the need for a revised naming prescription. Briefings in Functional Genomics 15: 65–71.

Chamberlain AJ, Vander Jagt CJ, Hayes BJ, Khansefid M, Marett LC, Millen CA, Nguyen TTT, Goddard ME. 2015. Extensive variation between tissues in allele specific expression in an outbred mammal. BMC Genomics 16: 993.

Chang H, Lim J, Ha M, Kim VN. 2014. TAIL-seq: Genome-wide Determination of Poly(A) Tail Length and 30 End Modifications. Molecular Cell 53:1044–1052.

Cheng J-H, Pan DZ-C, Tsai ZT-Y, Tsai H-K. 2015. Genome-wide analysis of enhancer RNA in gene regulation across 12 mouse tissues. Scientific Reports 5: 12648.

Chi SW, Zang JB, Mele A, Darnell RB. 2009. Argonaute HITS-CLIP decodes microRNA-mRNA interaction maps. Nature 460: 479.

Choo KB, Soon YL, Nguyen PNN, Hiew MSY, Huang C-J. 2014. MicroRNA-5p and -3p co-expression and cross-targeting in colon cancer cells. Journal of Biomedical Science 21: 95.

Chou C-H, Shrestha S, Yang C-D, Chang N-W, Lin Y-L, Liao K-W, Huang W-C, Sun T-H, Tu S-J, Lee W-H et al. 2018. miRTarBase update 2018: a resource for experimentally validated microRNA-target interactions. Nucleic Acids Research 46: D296–D302.

Clop A, Marcq F, Takeda H, Pirottin D, Tordoir X, Bibe B, Bouix J, Caiment F, Elsen J-M, Eychenne F, et al. 2006. A mutation creating a potential illegitimate microRNA target site in the myostatin gene affects muscularity in sheep. Nat Genet 38: 813–818.

Cowled C, Foo C-H, Deffrasnes C, Rootes CL, Williams DT, Middleton D, Wang L-F, Bean AGD, Stewart CR. 2017. Circulating microRNA profiles of Hendra virus infection in horses. Scientific Reports 7: 7431.

Creyghton MP, Cheng AW, Welstead GG, Kooistra T, Carey BW, Steine EJ, Hanna J, Lodato MA, Frampton GM, Sharp PA et al. 2010. Histone H3K27ac separates active from poised enhancers and predicts developmental state. Proceedings of the National Academy of Sciences of the United States of America 107: 21931–21936.

Daetwyler HD, Brauning R, Chamberlain AJ, McWilliam S, McCulloch A, Jagt CJV, Sunduimijid B, Hayes BJ, Kijas JW. 2017. 1000 Bull Genomes and SheepGenomeDB Projects: Enabling Cost-Effective Sequence Level Analyses Globally. In Association for the Advancement of Animal Breeding and Genetics, Rydges Southbank Townsville Queensland.

Daetwyler HD, Capitan A, Pausch H, Stothard P, van Binsbergen R, Brondum RF, Liao X, Djari A, Rodriguez SC, Grohs C et al. 2014. Whole-genome sequencing of 234 bulls facilitates mapping of monogenic and complex traits in cattle. Nat Genet 46: 858–865.

Dobin A, Davis CA, Schlesinger F, Drenkow J, Zaleski C, Jha S, Batut P, Chaisson M, Gingeras TR. 2013. STAR: ultrafast universal RNA-seq aligner. Bioinformatics 29: 15–21.

Dowen Jill M, Fan Zi P, Hnisz D, Ren G, Abraham Brian J, Zhang Lyndon N, Weintraub Abraham S, Schuijers J, Lee Tong I, Zhao K et al. 2014. Control of Cell Identity Genes Occurs in Insulated Neighborhoods in Mammalian Chromosomes. Cell 159: 374–387.

Dweep H, Gretz N. 2015. miRWalk2.0: a comprehensive atlas of microRNA-target interactions. Nature Methods 12: 697.

ENCODE Project Consortium. 2012. An Integrated Encyclopedia of DNA Elements in the Human Genome. Nature 489: 57–74.

Friedlander MR, Chen W, Adamidi C, Maaskola J, Einspanier R, Knespel S, Rajewsky N. 2008. Discovering microRNAs from deep sequencing data using miRDeep. Nature Biotechnology 26: 407.

Friedlander MR, Mackowiak SD, Li N, Chen W, Rajewsky N. 2012. miRDeep2 accurately identifies known and hundreds of novel microRNA genes in seven animal clades. Nucleic Acids Research 40: 37–52.

Friedman RC, Farh KK, Burge CB, Bartel DP. 2009. Most mammalian mRNAs are conserved targets of microRNAs. Genome Res 19.

Gai Y, Zhang J, Wei C, Cao W, Cui Y, Cui S. 2017. miR-375 negatively regulates the synthesis and secretion of catecholamines by targeting Sp1 in rat adrenal medulla. American Journal of Physiology-Cell Physiology 312: C663–C672.

Garcia DM, Baek D, Shin C, Bell GW, Grimson A, Bartel DP. 2011. Weak seed-pairing stability and high target-site abundance decrease the proficiency of lsy-6 and other microRNAs. Nature Structural & Molecular Biology 18:1139.

Gong J, Tong Y, Zhang HM, Wang K, Hu T, Shan G, Sun J, Guo AY. 2012. Genome-wide identification of SNPs in microRNA genes and the SNP effects on microRNA target binding and biogenesis. Hum Mu tat 33.

Gong J, Wu Y, Zhang X, Liao Y, Sibanda VL, Liu W, Guo A-Y. 2014. Comprehensive analysis of human small RNA sequencing data provides insights into expression profiles and miRNA editing. RNA Biology 11: 1375–1385.

Griffiths-Jones S, Saini HK, Dongen S, Enright AJ. 2008. miRBase: tools for microRNA genomics. Nucleic Acids Res 36.

Griffiths-Jones S, Hui JHL, Marco A, Ronshaugen M. 2011. MicroRNA evolution by arm switching. EMBO reports 12: 172–177.

Grimson A, Farh KK-H, Johnston WK, Garrett-Engele P, Lim LP, Bartel DP. 2007. MicroRNA Targeting Specificity in Mammals: determinants beyond Seed Pairing. Mol Cell 27.

Grisart B, Farnir P, Karim L, Cambisano N, Kim JJ, Kvasz A, Mni M, Simon P, Frère JM, Coppieters W et al. 2004. Genetic and functional confirmation of the causality of the DGAT1 K232A quantitative trait nucleotide in affecting milk yield and composition. Proceedings of the National Academy of Sciences of the United States of America 101: 2398–2403.

Grosswendt S, Filipchyk A, Manzano M, Klironomos F, Schilling M, Herzog M, Gottwein E, Rajewsky N. 2014. Unambiguous Identification of miRNA:Target Site Interactions by Different Types of Ligation Reactions. Molecular Cell 54: 1042–1054.

Gu Z, Eleswarapu S, Jiang H. 2007. Identification and characterization of microRNAs from the bovine adipose tissue and mammary gland. FEBS Lett 581.

Guo Z, Maki M, Ding R, Yang Y, zhang B, Xiong L. 2014. Genome-wide survey of tissue-specific microRNA and transcription factor regulatory networks in 12 tissues. Scientific Reports 4: 5150.

Hafner M, Landthaler M, Burger L, Khorshid M, Hausser J, Berninger P, Rothballer A, Ascano M, Jungkamp AC, Munschauer M, et al. 2010. Transcriptome-wide identification of RNA-binding protein and microRNA target sites by PAR-CLIP. Cell 141.

Hasuwa H, Ueda J, Ikawa M, Okabe M. 2013. MiR-200b and miR-429 Function in Mouse Ovulation and Are Essential for Female Fertility. Science 341: 71–73.

Helwak A, Kudla G, Dudnakova T, Tollervey D. 2013. Mapping the Human miRNA Interactome by CLASH Reveals Frequent Noncanonical Binding. Cell 153: 654–665.

Hu W, Wang T, Yue E, Zheng S, Xu J-H. 2014. Flexible microRNA arm selection in rice. Biochemical and Biophysical Research Communications 447: 526–530.

Huang Y, Ren HT, Xiong JL, Gao XC, Sun XH. 2017. Identification and characterization of known and novel microRNAs in three tissues of Chinese giant salamander base on deep sequencing approach. Genomics 109: 258–264.

Iborra FJ, Pombo A, Jackson DA, Cook PR. 1996. Active RNA polymerases are localized within discrete transcription “factories’ in human nuclei. Journal of Cell Science 109: 1427–1436.

Ioannidis J, Donadeu FX. 2017. Changes in circulating microRNA levels can be identified as early as day 8 of pregnancy in cattle. PLOS ONE 12: e0174892.

Jan CH, Friedman RC, Ruby JG, Bartel DP. 2010. Formation, regulation and evolution of Caenorhabditis elegans 30UTRs. Nature 469: 97.

Ji Z, Liu Z, Chao T, Hou L, Fan R, He R, Wang G, Wang J. 2017. Screening of miRNA profiles and construction of regulation networks in early and late lactation of dairy goat mammary glands. Scientific Reports 7: 11933.

Jin W, Grant JR, Stothard P, Moore SS, Guan LL. 2009. Characterization of bovine miRNAs by sequencing and bioinformatics analysis. BMC Mol Biol 10.

Jopling C. 2012. Liver-specific microRNA-122: Biogenesis and function. RNA Biol 9.

Joshi NA, Fass JN. 2011. Sickle: A sliding-window, adaptive, quality-based trimming tool for FastQ files [Software]. Available at https://github.com/najoshi/sickle.

Khan A, Zhang X. 2016. dbSUPER: a database of super-enhancers in mouse and human genome. Nucleic Acids Research 44: D164–D171.

Kim D, Langmead B, Salzberg SL. 2015. HISAT: a fast spliced aligner with low memory requirements. Nature Methods 12: 357.

Kloosterman WP, Plasterk RHA. 2006. The Diverse Functions of MicroRNAs in Animal Development and Disease. Developmental Cell 11: 441–450.

Kodzius R, Kojima M, Nishiyori H, Nakamura M, Fukuda S, Tagami M, Sasaki D, Imamura K, Kai C, Harbers M et al. 2006. CAGE: cap analysis of gene expression. Nat Meth 3: 211–222.

Kozomara A, Griffiths-Jones S. 2011. miRBase: integrating microRNA annotation and deep­sequencing data. Nucleic Acids Res 39.

Kozomara A, Griffiths-Jones S 2014. miRBase: annotating high confidence microRNAs using deep sequencing data. Nucleic Acids Res 42.

Krek A, Grün D, Poy MN, Wolf R, Rosenberg L, Epstein EJ, MacMenamin P, da Piedade I, Gunsalus KC, Stoffel M et al. 2005. Combinatorial microRNA target predictions. Nat Genet 37.

Kren BT, Wong PYP, Sarver A, Zhang X, Zeng Y, Steer CJ. 2009. microRNAs identified in highly purified liver-derived mitochondria may play a role in apoptosis. RNA biology 6: 65–72.

Kuchenbauer F, Mah SM, Heuser M, McPherson A, Rüschmann J, Rouhi A, Berg T, Bullinger L, Argiropoulos B, Morin RD et al. 2011. Comprehensive analysis of mammalian miRNA* species and their role in myeloid cells. Blood 118: 3350–3358.

Kuhn RM, Haussler D, Kent WJ. 2013. The UCSC genome browser and associated tools. Briefings in Bioinformatics 14: 144–161.

Kuo W-T, Su M-W, Lee Y, Chen C-H, Wu C-W, Fang W-L, Huang K-H, Lin W-c. 2015. Bioinformatic Interrogation of 5p-arm and 3p-arm Specific miRNA Expression Using TCGA Datasets. Journal of Clinical Medicine 4: 1798.

Kwak H, Fuda NJ, Core LJ, Lis JT. 2013. Precise Maps of RNA Polymerase Reveal How Promoters Direct Initiation and Pausing. Science 339: 950–953.

Lagana A, Veneziano D, Spata T, Tang R, Zhu H, Mohler PJ, Kilic A. 2015. Identification of General and Heart-Specific miRNAs in Sheep (Ovis aries). PLOS ONE 10: e0143313.

Lai X, Vera J. 2013. MicroRNA Clusters. In Encyclopedia of Systems Biology, (ed. W Dubitzky, O Wolkenhauer, K-H Cho, H Yokota), pp. 1310–1314. Springer New York, New York, NY.

Langmead B. 2010. Aligning short sequencing reads with Bowtie. Curr Protoc Bioinformatics Chapter 11.

Langmead B, Salzberg SL. 2012. Fast gapped-read alignment with Bowtie 2. Nat Methods 9.

Lee RC, Feinbaum RL, Ambros V. 1993. The C. elegans heterochronic gene lin-4 encodes small RNAs with antisense complementarity to lin-14. Cell 75: 843–854.

Lee Y, Kim M, Han J, Yeom KH, Lee S, Baek SH, Kim VN. 2004. MicroRNA genes are transcribed by RNA polymerase II. The EMBO Journal 23: 4051–4060.

Leung AKL, Sharp PA. 2010. MicroRNA Functions in Stress Responses. Molecular Cell 40: 205–215.

Lewis BP, Burge CB, Bartel DP. 2005. Conserved Seed Pairing, Often Flanked by Adenosines, Indicates that Thousands of Human Genes are MicroRNA Targets. Cell 120: 15–20.

Li G, Wu X, Qian W, Cai H, Sun X, Zhang W, Tan S, Wu Z, Qian P, Ding K et al. 2016. CCAR1 50 UTR as a natural miRancer of miR-1254 overrides tamoxifen resistance. Cell Research 26: 655.

Li H, Durbin R. 2009. Fast and accurate short read alignment with Burrows-Wheeler transform. Bioinformatics 25: 1754–1760.

Li H, Handsaker B, Wysoker A, Fennell T, Ruan J, Homer N, Marth G, Abecasis G, Durbin R. 2009. The Sequence Alignment/Map format and SAMtools. Bioinformatics 25: 2078–2079.

Li H, Homer N. 2010. A survey of sequence alignment algorithms for next-generation sequencing. Briefings in Bioinformatics 11: 473–483.

Li M, Belmonte JCI. 2015. Roles for noncoding RNAs in cell-fate determination and regeneration. Nature Structural& Molecular Biology 22: 2.

Li S-C, Liao Y-L, Chan W-C, Ho M-R, Tsai K-W, Hu L-Y, Lai C-H, Hsu C-N, Lin W-c. 2011. Interrogation of rabbit miRNAs and their isomiRs. Genomics 98: 453–459.

Li S-C, Tsai K-W, Pan H-W, Jeng Y-M, Ho M-R, Li W-H. 2012. MicroRNA 3’ end nucleotide modification patterns and arm selection preference in liver tissues. BMC Systems Biology 6: S14.

Licatalosi DD, Mele A, Fak JJ, Ule J, Kayikci M, Chi SW, Clark TA, Schweitzer AC, Blume JE, Wang X, et al. 2008. HITS-CLIP yields genome-wide insights into brain alternative RNA processing. Nature 456.

Lim LP, Lau NC, Garrett-Engele P, Grimson A, Schelter JM, Castle J, Bartel DP, Linsley PS, Johnson JM. 2005. Microarray analysis shows that some microRNAs downregulate large numbers of target mRNAs. Nature 433: 769.

Lin M-H, Chen Y-Z, Lee M-Y, Weng K-P, Chang H-T, Yu S-Y, Dong B-J, Kuo F-R, Hung L-T, Liu L-F et al. 2018. Comprehensive identification of microRNA arm selection preference in lung cancer: miR-324-5p and -3p serve oncogenic functions in lung cancer. Oncology Letters 15: 9818­9826.

Londin E, Loher P, Telonis AG, Quann K, Clark P, Jing Y, Hatzimichael E, Kirino Y, Honda S, Lally M et al. 2015. Analysis of 13 cell types reveals evidence for the expression of numerous novel primate- and tissue-specific microRNAs. Proceedings of the National Academy of Sciences 112: E1106–E1115.

Lorenz R, Bernhart SH, Höner zu Siederdissen C, Tafer H, Flamm C, Stadler PF, Hofacker IL. 2011. ViennaRNA Package 2.0. Algorithms for Molecular Biology 6: 26.

Love Ml, Huber W, Anders S. 2014. Moderated estimation of fold change and dispersion for RNA-seq data with DESeq2. Genome Biology 15: 550.

Ludwig N, Leidinger P, Becker K, Backes C, Fehlmann T, Pallasch C, Rheinheimer S, Meder B, Stahler C, Meese E et al. 2016. Distribution of miRNA expression across human tissues. Nucleic Acids Research 44: 3865–3877.

Lutzow YCS, Donaldson L, Gray CP, Vuocolo T, Pearson RD, Reverter A, Byrne KA, Sheehy PA, Windon R, Tellam RL. 2008. Identification of immune genes and proteins involved in the response of bovine mammary tissue to Staphylococcus aureus infection. BMC Veterinary Research 4: 18.

Magee RG, Telonis AG, Loher P, Londin E, Rigoutsos I. 2018. Profiles of miRNA Isoforms and tRNA Fragments in Prostate Cancer. Scientific Reports 8: 5314.

Mallory AC, Reinhart BJ, Jones-Rhoades MW, Tang G, Zamore PD, Barton MK, Bartel DP. 2004. MicroRNA control of *PHABULOSA* in leaf development: importance of pairing to the microRNA 50 region. The EMBO Journal 23: 3356–3364.

Marco A, Hui JHL, Ronshaugen M, Griffiths-Jones S. 2010. Functional Shifts in Insect microRNA Evolution. Genome Biology and Evolution 2: 686–696.

Marco A, Ninova M, Ronshaugen M, Griffiths-Jones S. 2013. Clusters of microRNAs emerge by new hairpins in existing transcripts. Nucleic Acids Research 41: 7745–7752.

Marson A, Levine SS, Cole MF, Frampton GM, Brambrink T, Johnstone S, Guenther MG, Johnston WK, Wernig M, Newman J et al. 2008. Connecting microRNA Genes to the Core Transcriptional Regulatory Circuitry of Embryonic Stem Cells. Cell 134: 521–533.

Martin M. 2011. Cutadapt removes adapter sequences from high-throughput sequencing reads. EMB Net J 17.

Mehta A, Baltimore D. 2016. MicroRNAs as regulatory elements in immune system logic. Nature Reviews Immunology 16: 279.

Melamed Ze, Levy A, Ashwal-Fluss R, Lev-Maor G, Mekahel K, Atias N, Gilad S, Sharan R, Levy C, Kadener S et al. 2013. Alternative Splicing Regulates Biogenesis of miRNAs Located across Exon-Intron Junctions. Molecular Cell 50: 869–881.

Meng Y, Shao C, Ma X, Wang H. 2013. Introns targeted by plant microRNAs: a possible novel mechanism of gene regulation. Rice 6: 8.

Mills RE, Luttig CT, Larkins CE, Beauchamp A, Tsui C, Pittard WS, Devine SE. 2006. An initial map of insertion and deletion (INDEL) variation in the human genome. Genome Research 16: 1182–1190.

Modepalli V, Kumar A, Hinds LA, Sharp JA, Nicholas KR, Lefevre C. 2014. Differential temporal expression of milk miRNA during the lactation cycle of the marsupial tammar wallaby (Macropus eugenii). BMC Genomics 15: 1012.

Moore SG, Pryce JE, Hayes BJ, Chamberlain AJ, Kemper KE, Berry DP, McCabe M, Cormican P, Lonergan P, Fair T et al. 2016. Differentially Expressed Genes in Endometrium and Corpus Luteum of Holstein Cows Selected for High and Low Fertility Are Enriched for Sequence Variants Associated with Fertility1. Biology of Reproduction 94: 19,11-11-19,11-11.

Morlando M, Ballarino M, Gromak N, Pagano F, Bozzoni I, Proudfoot NJ. 2008. Primary microRNA transcripts are processed co-transcriptionally. Nature Structural & Molecular Biology 15: 902.

Nojima T, Gomes T, Grosso Ana Rita F, Kimura H, Dye Michael J, Dhir S, Carmo-Fonseca M, Proudfoot Nicholas J. 2015. Mammalian NET-Seq Reveals Genome-wide Nascent Transcription Coupled to RNA Processing. Cell 161: 526–540.

Osborne CS, Chakalova L, Brown KE, Carter D, Horton A, Debrand E. 2004. Active genes dynamically colocalize to shared sites of ongoing transcription. Nat Genet 36.

Paicu C, Mohorianu I, Stocks M, Xu P, Coince A, Billmeier M, Dalmay T, Moulton V, Moxon S. 2017. miRCat2: accurate prediction of plant and animal microRNAs from next-generation sequencing datasets. Bioinformatics 33: 2446–2454.

Pawlicki JM, Steitz JA. 2008. Primary microRNA transcript retention at sites of transcription leads to enhanced microRNA production. The Journal of Cell Biology 182: 61–76.

Penso-Dolfin L, Swofford R, Johnson J, Alföldi J, Lindblad-Toh K, Swarbreck D, Moxon S, Di Palma F. 2016. An Improved microRNA Annotation of the Canine Genome. PLOS ONE 11: e0153453.

Place RF, Li L-C, Pookot D, Noonan EJ, Dahiya R. 2008. MicroRNA-373 induces expression of genes with complementary promoter sequences. Proceedings of the National Academy of Sciences 105: 1608–1613.

Reczko M, Maragkakis M, Alexiou P, Grosse I, Hatzigeorgiou AG. 2012. Functional microRNA targets in protein coding sequences. Bioinformatics 28: 771–776.

Reinhart BJ, Slack FJ, Basson M, Pasquinelli AE, Bettinger JC, Rougvie AE, Horvitz HR, Ruvkun G. 2000. The 21-nucleotide let-7 RNA regulates developmental timing in Caenorhabditis elegans. Nature 403: 901.

Renthal NE, Williams KrC, Mendelson CR. 2013. MicroRNAs—mediators of myometrial contractility during pregnancy and labour. Nature Reviews Endocrinology 9: 391.

Saunders MA, Liang H, Li W-H. 2007. Human polymorphism at microRNAs and microRNA target sites. Proceedings of the National Academy of Sciences 104: 3300–3305.

Scheel TKH, Moore MJ, Luna JM, Nishiuchi E, Fak J, Darnell RB, Rice CM. 2017. Global mapping of miRNA-target interactions in cattle (Bos taurus). Scientific Reports 7: 8190.

Schnall-Levin M, Zhao Y, Perrimon N, Berger B. 2010. Conserved microRNA targeting in *Drosophila* is as widespread in coding regions as in 30UTRs. Proceedings of the National Academy of Sciences 107: 15751–15756.

Selbach M, Schwanhäusser B, Thierfelder N, Fang Z, Khanin R, Rajewsky N. 2008. Widespread changes in protein synthesis induced by microRNAs. Nature 455: 58.

Sengar GS, Deb R, Singh U, Raja TV, Kant R, Sajjanar B, Alex R, Alyethodi RR, Kumar A, Kumar S et al. 2018. Differential expression of microRNAs associated with thermal stress in Frieswal (Bos taurus x Bos indicus) crossbred dairy cattle. Cell Stress and Chaperones 23: 155–170.

spicuglia s, Vanhille L. 2012. Chromatin signatures of active enhancers. Nucleus 3: 126–131.

Stark A, Bushati N, Jan CH, Kheradpour P, Hodges E, Brennecke J, Bartel DP, Cohen SM, Kellis M. 2008. A single Hox locus in Drosophila produces functional microRNAs from opposite DNA strands. Genes & Development 22: 8–13.

Stark A, Lin MF, Kheradpour P, Pedersen JS, Parts L, Carlson JW, Crosby MA, Rasmussen MD, Roy S, Deoras AN et al. 2007. Discovery of functional elements in 12 Drosophila genomes using evolutionary signatures. Nature 450: 219.

Suzuki HI, Young RA, Sharp PA. 2017. Super-Enhancer-Mediated RNA Processing Revealed by Integrative MicroRNA Network Analysis. Cell 168: 1000–1014.el015.

Szabo G, Bala S. 2013. MicroRNAs in liver disease. Nature Reviews Gastroenterology & Hepatology 10: 542.

Telonis AG, Loher P, Jing Y, Londin E, Rigoutsos I. 2015. Beyond the one-locus-one-miRNA paradigm: microRNA isoforms enable deeper insights into breast cancer heterogeneity. Nucleic Acids Research 43: 9158–9175.

Tilgner H, Jahanbani F, Blauwkamp T, Moshrefi A, Jaeger E, Chen F. 2015. Comprehensive transcriptome analysis using synthetic long-read sequencing reveals molecular co­association of distant splicing events. Nat Biotechnol 33.

Tsai K-W, Leung C-M, Lo Y-H, Chen T-W, Chan W-C, Yu S-Y, Tu Y-T, Lam H-C, Li S-C, Ger L-P et al. 2016. Arm Selection Preference of MicroRNA-193a Varies in Breast Cancer. Scientific Reports 6: 28176.

Tyler DM, Okamura K, Chung W-J, Hagen JW, Berezikov E, Hannon GJ, Lai EC. 2008. Functionally distinct regulatory RNAs generated by bidirectional transcription and processing of microRNA loci. Genes & Development 22: 26–36.

Villar D, Berthelot C, Aldridge S, Rayner Tim F, Lukk M, Pignatelli M, Park Thomas J, Deaville R, Erichsen Jonathan T, Jasinska Anna J et al. 2015. Enhancer Evolution across 20 Mammalian Species. Cell 160: 554–566.

Wake C, Labadorf A, Dumitriu A, Hoss AG, Bregu J, Albrecht KH, DeStefano AL, Myers RH. 2016. Novel microRNA discovery using small RNA sequencing in post-mortem human brain. BMC Genomics 17: 776.

Wang D, Liang G, Wang B, Sun H, Liu J, Guan LL. 2016a. Systematic microRNAome profiling reveals the roles of microRNAs in milk protein metabolism and quality: insights on low-quality forage utilization. Scientific Reports 6: 21194.

Wang M, Hancock TP, MacLeod IM, Pryce JE, Cocks BG, Hayes BJ. 2017. Putative enhancer sites in the bovine genome are enriched with variants affecting complex traits. Genetics Selection Evolution 49: 56.

Wang W, Kwon EJ, Tsai L-H. 2012. MicroRNAs in learning, memory, and neurological diseases. Learning & Memory 19: 359–368.

Wang Y, Luo J, Zhang H, Lu J. 2016b. microRNAs in the Same Clusters Evolve to Coordinately Regulate Functionally Related Genes. Molecular Biology and Evolution 33: 2232–2247.

Whyte WA, Orlando DA, Hnisz D, Abraham BJ, Lin CY, Kagey MH. 2013. Master transcription factors and mediator establish super-enhancers at key cell identity genes. Cell 153.

Wong LL, Rademaker MT, Saw EL, Lew KS, Ellmers LJ, Charles CJ, Richards AM, Wang P. 2017. Identification of novel microRNAs in the sheep heart and their regulation in heart failure. Scientific Reports 7: 8250.

Xiao M, Li J, Li W, Wang Y, Wu F, Xi Y, Zhang L, Ding C, Luo H, Li Y et al. 2016. MicroRNAs activate gene transcription epigenetically as an enhancer trigger. RNA Biology: 1–9.

Yu J, Wang F, Yang G-H, Wang F-L, Ma Y-N, Du Z-W, Zhang J-W. 2006. Human microRNA clusters: Genomic organization and expression profile in leukemia cell lines. Biochemical and Biophysical Research Communications 349: 59–68.

Zhang Y, Fan M, Zhang X, Huang F, Wu K, Zhang J, Liu J, Huang Z, Luo H, Tao L et al. 2014. Cellular microRNAs up-regulate transcription via interaction with promoter TATA-box motifs. RNA 20: 1878–1889.

Zhao C, Carrillo JA, Tian F, Zan L, Updike SM, Zhao K, Zhan F, Song J. 2015. Genome-Wide H3K4me3 Analysis in Angus Cattle with Divergent Tenderness. PLoS One 10: e0115358.

Zhao Y, Samal E, Srivastava D. 2005. Serum response factor regulates a muscle-specific microRNA that targets Hand2 during cardiogenesis. Nature 436: 214.

Zhixiong L, Hongliang W, Ling C, Lijun W, Xiaolin L, Caixia R, Ailong S. 2014. Identification and characterization of novel and differentially expressed microRNAs in peripheral blood from healthy and mastitis Holstein cattle by deep sequencing. Animal Genetics 45: 20–27.

Zisoulis DG, Lovci MT, Wilbert ML, Hutt KR, Liang TY, Pasquinelli AE, Yeo GW. 2010. Comprehensive discovery of endogenous Argonaute binding sites in Caenorhabditis elegans. Nature Structural & Molecular Biology 17: 173.

