## Supplementary figures and images for "Genome-wide profiling of microRNAs and prediction of mRNA targets in 17 bovine tissues"

### Supplemental_Fig_S1.png

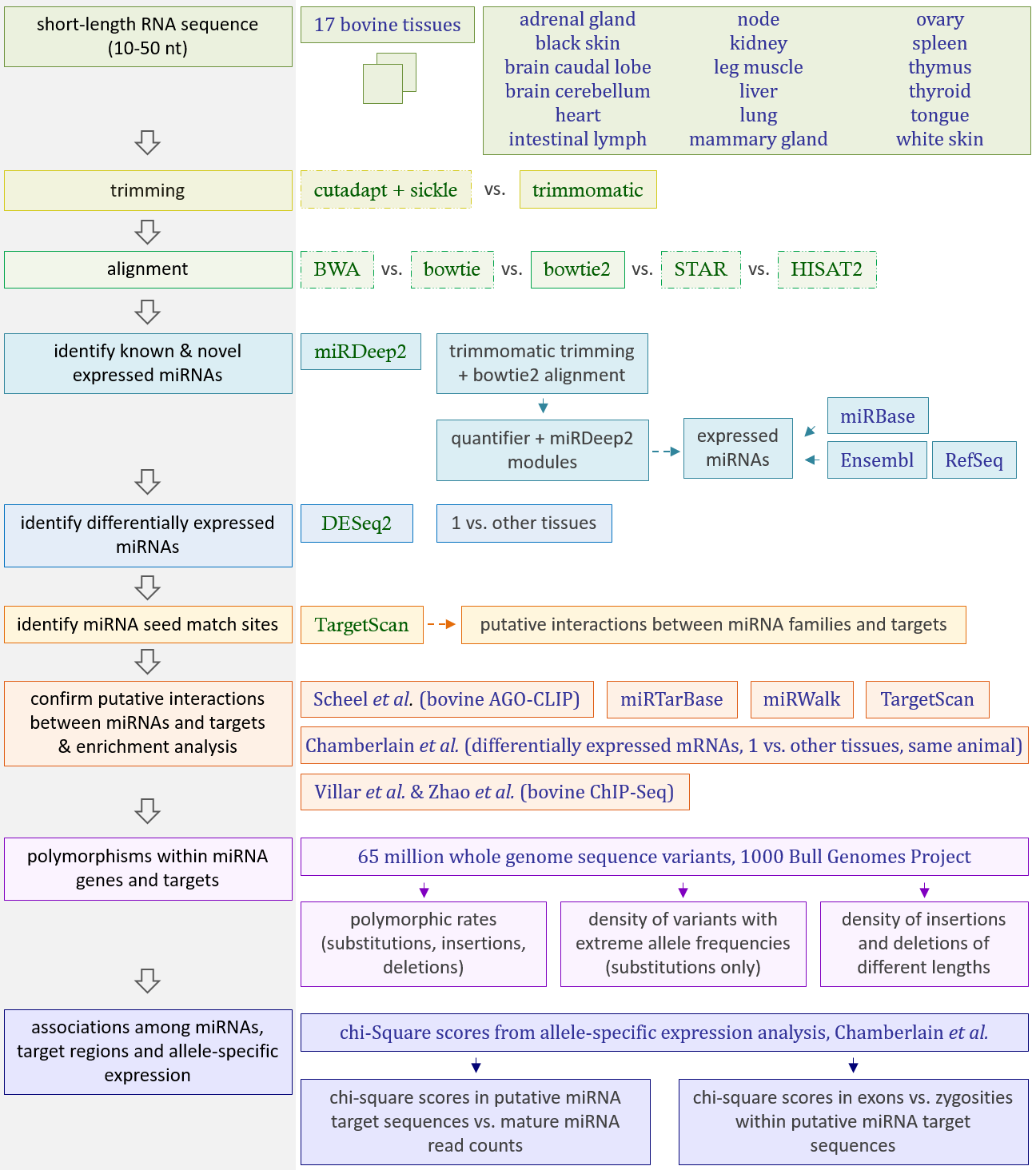
